# Beyond codon optimality: codon pairs regulate mRNA stability dependent on translation

**DOI:** 10.64898/2026.09.15.751551

**Authors:** Haejeong Lee, Damir Musaev, Charles E. Vejnar, Srikar Krishna, Trung Duc Nguyen, Ethan C. Strayer, Mario Abdel Messih, Carter M. Takacs, Jean-Denis Beaudoin, Antonio J. Giraldez

## Abstract

The coding sequence of an mRNA directs its own decay, yet how codons, codon context, and amino acids collectively regulate mRNA stability remains poorly understood. Here we use a massively parallel reporter assay to decode this regulatory layer in zebrafish embryos. We find that codon pairs and amino acid pairs regulate mRNA stability beyond the level of individual codons. This regulation depends on the identity and order of neighboring codons and their encoded amino acids in a translation-dependent manner. The relative contributions of codon and amino acid combinations to mRNA stability can be quantified using machine learning. We further found that endogenous mRNAs are regulated by codon pairs, thereby governing developmental gene regulation and biological function. This codon context-dependent decay requires deadenylation and decapping by Cnot7 and Dcp2, with Upf1 acting on long non-optimal ORFs. Together, our results redefine codon optimality as a context-dependent, pair-level code with implications for RNA biology and therapeutic mRNA design.

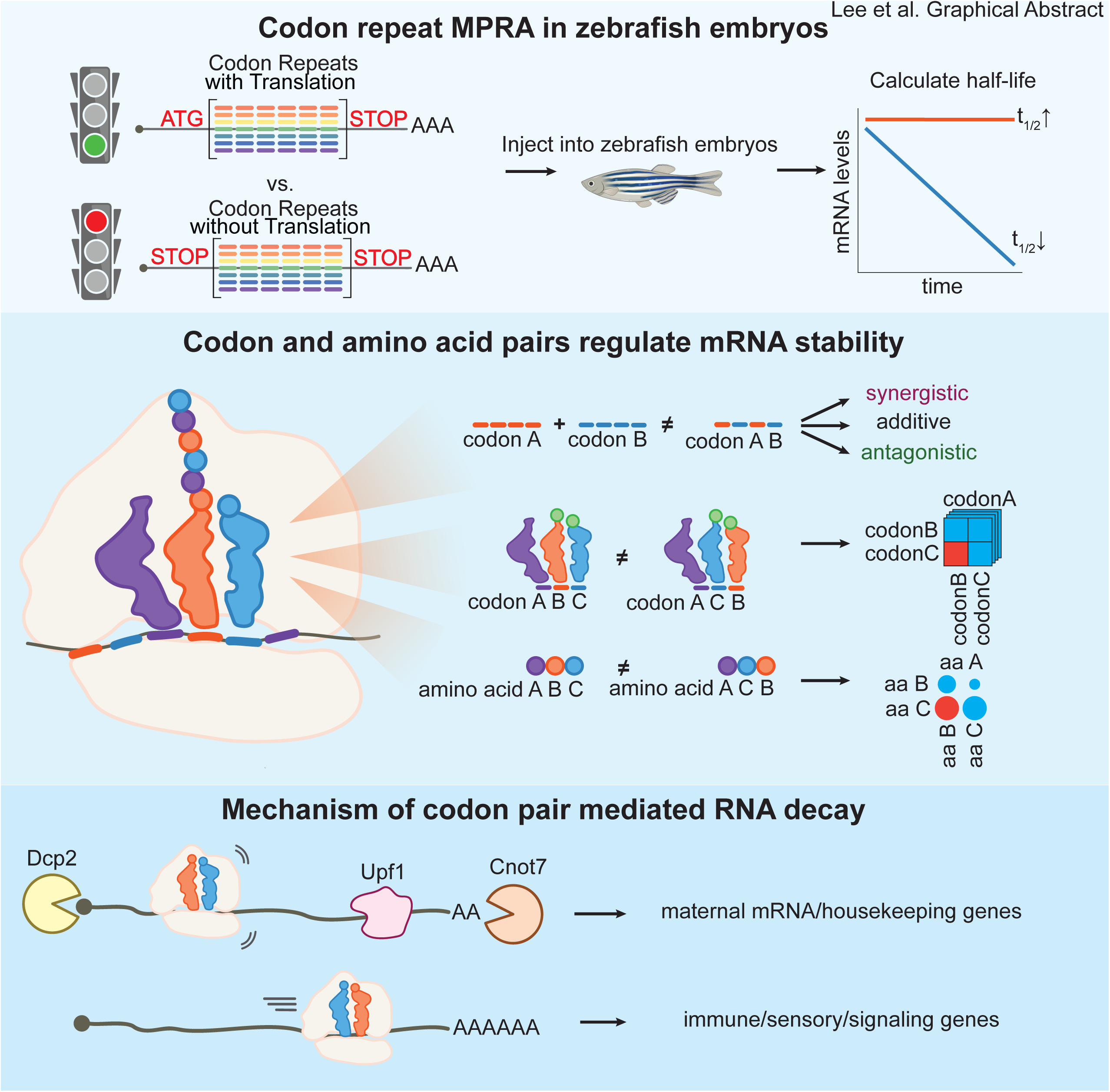

**Highlights:**

- Codon pairs act non-additively, creating synergistic and antagonistic mRNA decay
- mRNA stability depends on the identity and the order of codon and amino acid pairs
- Dcp2 and Cnot7 drive codon pair decay, with Upf1 working on long non-optimal ORFs
- Codon pair usage shapes maternal mRNA clearance and tracks gene function

## Introduction

mRNA stability is a fundamental regulator of gene expression across all domains of life ^1^, controlling genes central to proliferation, differentiation, and stress responses through regulated mRNA decay ^2^. mRNA stability also shapes multiple cellular and developmental transitions by facilitating the clearance of transcripts from one state to establish the next. Specifically, during the maternal-to-zygotic transition (MZT), a universal step in animal development, maternally deposited mRNAs are cleared after zygotic genome activation ^3^. Failure to do so causes embryogenesis defects in Drosophila, zebrafish, Xenopus, and humans ^4–7^, underscoring the importance of mRNA decay during development. Maternal mRNAs are regulated by multiple mechanisms encoded in the untranslated region (UTR), including AU-rich elements, microRNA binding sites, and 3’UTR length, among others ^4,5,7,8^. Additionally, clearance of these maternal mRNAs is translation-dependent degradation through codon optimality and CDS length ^9–11^.

mRNA stability is closely linked to translation through initiation efficiency, elongation rate, and ribosome-dependent decay ^1^. The CDS contributes to this relationship primarily through codon usage. The codon stability coefficient (CSC) quantifies this link by measuring the correlation between codon frequency and transcript half-life ^12^, with stabilizing codons having positive CSC values and destabilizing codons negative CSC values. The CSC value correlates with tRNA level, indicating codons paired with abundant tRNAs are decoded at faster rates and stabilize mRNA, whereas those decoded by rare tRNAs slow elongation and trigger decay ^9,12^. However, the correlation between tRNA abundance and codon optimality is only moderate, indicating that additional factors regulate codon-mediated stability. In yeast, ribosome stalling causes prolonged A-site dwell time detected by Dhh1p and Not5 of the CCR4-NOT deadenylase complex ^13–15^. More recent work in human cells suggests a more complex recognition of slow elongation by CNOT3 and DHX29 via separate mechanisms ^16,17^.

Growing evidence indicates that individual codon identity alone cannot fully account for CDS-mediated stability regulation ^18^. Analyses from diverse genomes suggest that codon pair usage is subject to evolutionary constraints beyond single-codon preferences ^19–21^. In yeast, 17 codon pairs strongly inhibit translation, with 12 showing codon order specificity ^22^; however, whether these effects operate in vertebrates and shape mRNA stability remains unknown. At the amino acid level, synonymous codons encoding the same amino acid often exert similar effects on mRNA half-life, implicating the nascent peptide in stability regulation ^9,23^. In addition, combinations of positively charged and bulky hydrophobic amino acids destabilize mRNAs through ribosome exit tunnel interactions, particularly when forming extended β-strand features ^24,25^. Despite these reports ^9,22–25^, a comprehensive and systematic characterization of how codons, codon pairs, amino acids, and amino acid pairs each contribute to mRNA stability, and the mechanisms behind them, remains lacking.

Progress has been limited by the complexity of endogenous transcripts, where confounding factors such as UTR elements, CDS length, and codon context make it difficult to isolate individual contributions. To overcome these challenges, we developed a reductionist approach inspired by the original simplified coding sequence experiments that deciphered the genetic code ^26^. We designed a comprehensive set of repetitive codon reporters to decode codon-optimality, thereby isolating the effects of individual elements and their combinations in a controlled manner. Here, we tested codon repeat massively parallel reporter libraries in zebrafish embryos and analyzed stability in both translation-dependent and translation-independent manners. We found that codon pairs exert stability effects distinct from the simple average of their constituent codons, and that these effects depend on the codon order. The resulting comprehensive codon-pair stability dataset enabled optimization of mRNA stability. We show that CDS-mediated stability is translation-dependent, with specific exceptions involving RNA secondary structure. We further identified amino acid pair properties and their order as predictors of stability effects. We dissected their contributions using machine learning, finding that the encoded amino acids explain the most variance while codon and codon pair usage add independent signals. In addition, we demonstrated that codon pairs regulate maternal mRNA clearance and couple transcript stability to gene function. Finally, we show that codon context-dependent decay requires the core decay factors Dcp2 and Cnot7, as well as the ORF-mediated decay (OMD) factor Upf1, but not the no-go decay factors Znf598 and Ski7. Taken together, our results demonstrate that mRNA stability is encoded within the CDS at multiple levels through the identity and order of codon pairs, and amino acid pairs.

## Results

### Neighboring codons determine mRNA stability beyond single codon optimality

Codon-mediated mRNA stability has been attributed to individual codon identity and whether a codon is decoded by abundant or rare tRNAs ^9,12^. However, the complexity of endogenous sequences can mask higher order regulatory interactions beyond individual codons. To address this, we developed a massively parallel reporter assay (MPRA) with simple codon repeats that enable systematic half-life measurements for all 61 individual codons, 1,830 codon pairs, and 71,980 codon triplets in developing zebrafish embryos (Fig. 1A). These synthetic reporters share common 5’ and 3’ UTR sequences with a fixed poly(A) tail, thus isolating differences in mRNA stability to the coding sequence alone. We injected mRNA encoding these libraries at the 1-cell stage and measured their half-lives from 3 to 8 hours post-fertilization (hpf). This allows us to measure reporter decay directly without metabolic labeling or transcriptional inhibition (Fig. 1B, S1A). The MPRA reporter half-lives correlated with the previously defined CSC values derived from endogenous transcripts and with cognate tRNA abundances ^9,27^ (Spearman’s R 0.469 and 0.220, respectively; Fig. 1C-D, Fig. S1B-J), indicating that our MPRA recapitulates codon-mediated regulation while maintaining a simple codon organization that allows us to dissect higher order interactions.

**Figure 1.**
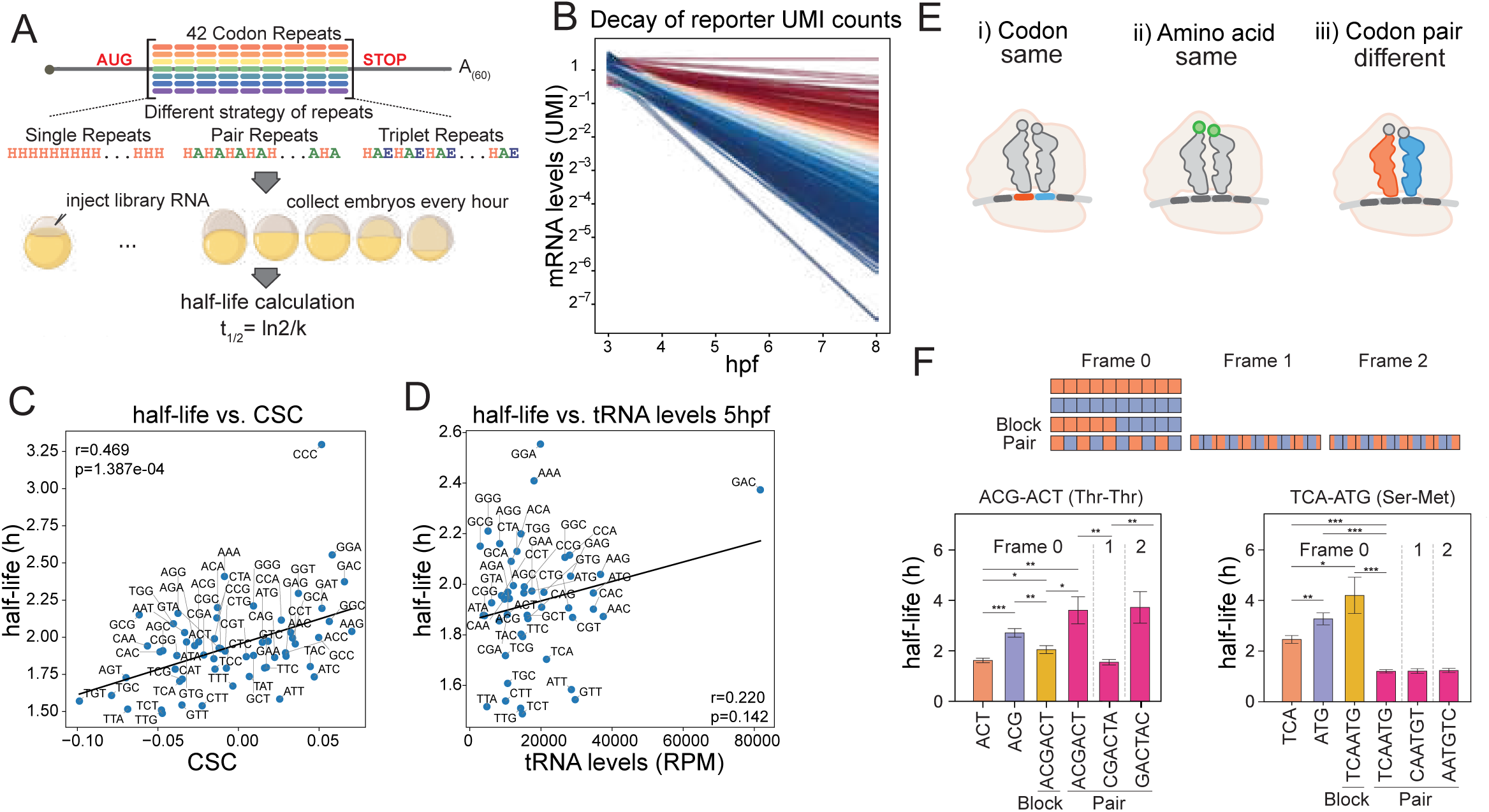
Codon pairs regulate stability beyond individual codons. **(A)** Schematics showing the design of the MPRA with repeats of single, pair, and triplet codons; mRNAs were injected into zebrafish embryos, collected every hour, and half-life was calculated. **(B)** Time-course decay of reporter UMI counts fitted to first-order exponential decay. Trendline generated from three biological replicates per hour from 3hpf to 8hpf. **(C and D)** Scatter plot comparing each reporter’s half-life to CSC (Codon Stability Coefficient) (C) and tRNA levels in a 5 hpf zebrafish embryo (D) (Spearman’s R). See also Figure S1. **(E)** Schematics comparing reporter sets with the same codon and amino acid composition but different codon pairs. **(F)** Schematics of reporters, with each codon shown as a square box; note that block and pair reporters share identical codon composition (top). Half-life comparison of reporters carrying two single-codon repeats, two codon mixtures in block versus pair arrangements, and pair reporters in different reading frames (bottom). Note that the pair reporters have significantly higher and lower half-lives than the block reporter, and that their half-lives are frame-dependent, suggesting translation dependency. Error bars show standard error, and we performed Welch’s t-test: ∗p < 0.05, ∗∗p < 0.01, ∗∗∗p < 0.001. See also Figure S2.

We hypothesized that if codon identity alone determines CDS-mediated mRNA stability, then stability should be independent of codon order. Conversely, if neighboring codons contribute to stability, rearranging codon order while preserving codon composition should alter mRNA half-life. To test this hypothesis, we first compared the stability of reporter mRNAs with single codon repeats (A_42_), codon pairs ([AB]_21_), and alternating groups of 5 identical codons ([5A5B]_4_[AA], blocks). We identified several lines of evidence indicating that codon pairs are a widespread determinant of mRNA stability. First, we observed that some codon pairs increase while others decrease mRNA half-life compared to the effect of individual codons. For example, comparing the half-lives of reporters encoding threonine homopolymer peptides with the same codon composition (e.g., ACG and ACT) but different codon arrangements (Fig. 1E) showed that alternating ACG and ACT codons conferred 1.7-fold higher stability than the alternating-block reporters, revealing a synergistic effect that depends on codon adjacency rather than composition alone (Fig. 1F). Conversely, alternating codons of TCA (Ser) and ATG (Met) conferred 3.5-fold lower stability than alternating-block reporters, indicating an antagonistic neighbor effect for these codons (Fig. 1F). Extending this analysis across 334 pairs revealed that 57% of codon pairs (190) have significantly different half-lives when arranged in blocks vs. individual pairs (Fig. S2), demonstrating that codon pair identity is a widespread determinant of stability. Second, the stability of the alternating-codon pair reporter depended on reading frame (Fig. 1F, S3A-C), suggesting that the codon context-driven mRNA stability regulation is linked to translation rather than to a sequence motif or RNA structure. Third, the effect of a codon on mRNA stability is influenced by its neighboring codon, resulting in synergistic, antagonistic, or additive effects (Fig. 2A, C-D). Systematic comparison of single codons versus the codon pairs across 1,501 codon pairs (Fig. 2C) identified 58 synergistic and 165 antagonistic codon pairs that exert significantly different stability compared to the average half-life of their constituent single codons (Benjamini-Hochberg-adjusted p<0.05; Fig. 2D-E). For example, ACG yields very different stabilities depending on its neighbor, destabilizing RNA when adjacent to ACA (half-life 2.21h) but stabilizing it when adjacent to ACT (half-life 3.61h), despite ACA and ACT having similar individual half-lives (1.66h and 1.76h, respectively) (Fig. 2B). Notably, specific codons, including GGA (Gly), CTC (Leu), and TCC (Ser), were particularly biased toward antagonistic interactions with other codons, despite being stabilizing or neutral as individual codons (Fig. 2D). Together, these results indicate that the codon pair context can influence the intrinsic stability of individual codons, revealing a layer of regulation that single codons alone fail to capture.

**Figure 2.**
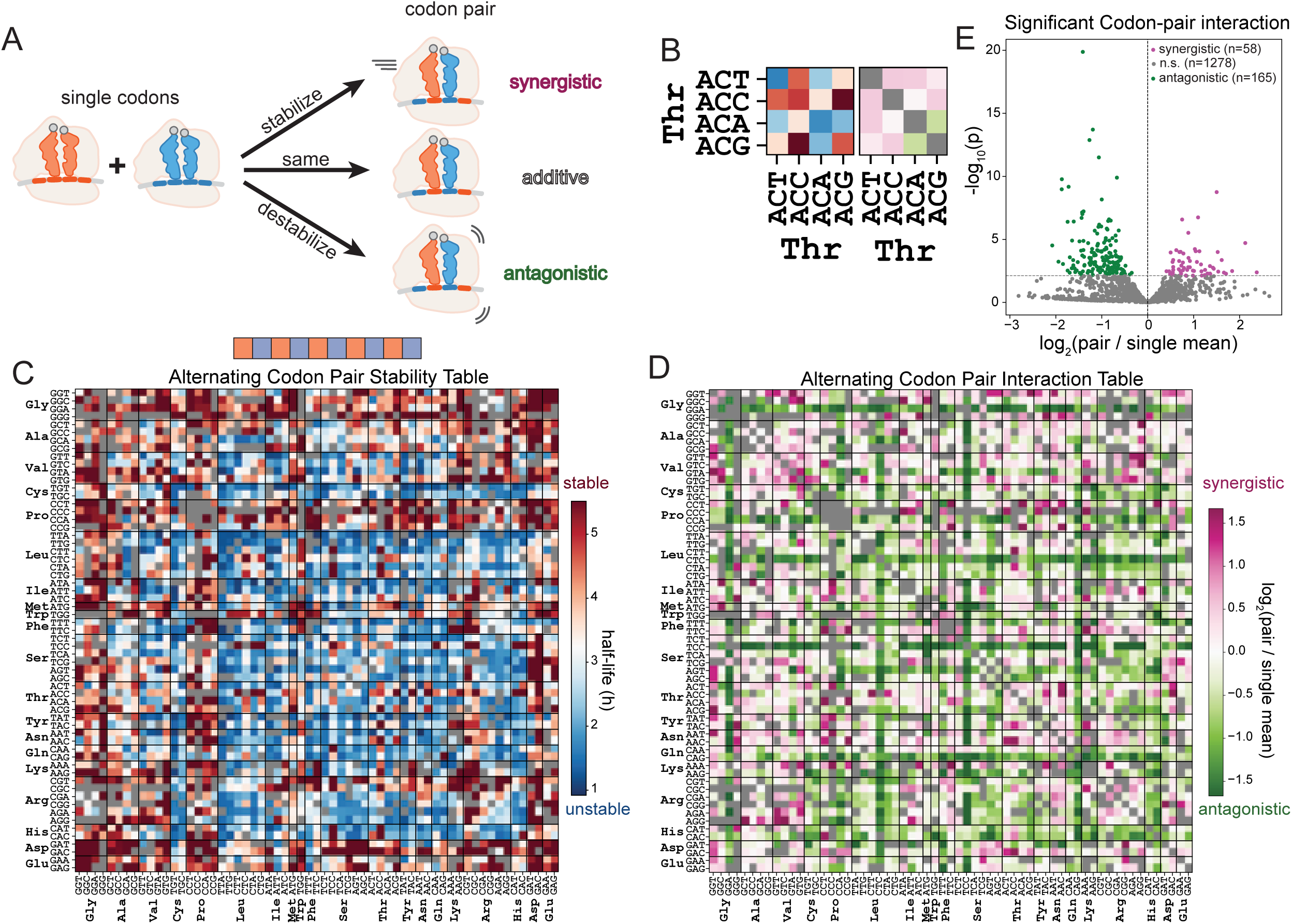
Codon pair stability is synergistic, additive, and antagonistic. **(A)** Schematics showing RNA half-life of codon pairs compared to single codons can be classified into synergistic, additive, and antagonistic. **(B)** Representative Threonine-Threonine codon pair combinations in RNA half-life heatmap (left; red, stable; blue, unstable) and codon pair interaction heatmap representing log_2_(pair half-life/mean of single half-lives) (right; pink, synergistic; green, antagonistic; gray, not applicable or no data). **(C)** Heatmap representing half-life of all alternating codon pair combinations (red, stable; blue, unstable; gray, no data). **(D)** Heatmap representing codon pair half-life change from the constituting codons log_2_(pair half-life/mean of single half-lives) (pink, synergistic; white, additive; green, antagonistic; gray in diagonal, not applicable; gray, no data). Note that the patterns of stability and codon pair interaction are distinct. **(E)** Volcano plot representing codon pair interaction score (log_2_(pair half-life / mean of single half-lives)) with its statistical significance (-log_10_(p)). We performed a two-sided z-test per codon pair, with the score’s standard error propagated from the half-life measurement and corrected with Benjamini-Hochberg FDR correction (pink, FDR < 0.05 and synergistic; green, FDR < 0.05 and antagonistic; gray, not significant).

### Codon pair effects depend on the order

Having established that codon pairs regulate mRNA stability, we next asked whether this effect depends on the order in which adjacent codons are decoded. If codon order is a determinant of stability, codon pair AB and BA should confer distinct half-lives. To test this, we designed 71,980 triplet-codon reporters in which the same three codons appear in two arrangements (ABC or ACB), reversing each internal pair (AB vs. BA, BC vs. CB, CA vs. AC) while holding codon composition fixed (Fig. 3A). Averaging half-lives across all 61 codon contexts for all triplets (NBC and NCB) allows comparion of the half-life difference between BC and CB, revealing how the order of each codon pair determines mRNA stability (Fig. 3B–D, Movie S1). Of 3,566 codon pairs, 44.5% (1,588) showed significantly different half-lives (BC vs. CB), indicating that codon-pair order is a prevalent determinant of mRNA stability (Fig. 3D). Resolving these order-dependent effects for each codon across all 61 x 61 codon neighbor combinations (ANN), reveals how the codon’s stability contribution depends on its neighbors, from the stabilizing GGG to the destabilizing TCT (Fig. 3E-H). These findings demonstrate that single-codon metrics such as CSC fail to account for the regulatory contribution of neighboring codons (Fig. 3I). To test whether codon-pair order regulates mRNA stability in other sequences beyond the repeat MPRAs, we inserted six codon triplets with identical composition but reversed order (ABC vs. ACB) into a GFP reporter. We observed that codon order regulates mRNA levels, despite identical codon content, recapitulating the MPRA results (Fig. 3J). Finally, to test whether this order dependence can be exploited to modulate mRNA stability, we used dynamic programming on codon triplet half-life to recode luciferase with an identical amino acid sequence but with stabilizing or destabilizing codon pairs. The stabilizing variant was 50% more stable than the destabilizing one (Fig. 3K), establishing that codon-pair order is not only a regulatory principle but a programmable one. Together, these results indicate that codon-pair order, not just codon composition, determines mRNA stability, revealing a layer of CDS-encoded regulation beyond the resolution of single-codon optimality.

**Figure 3.**
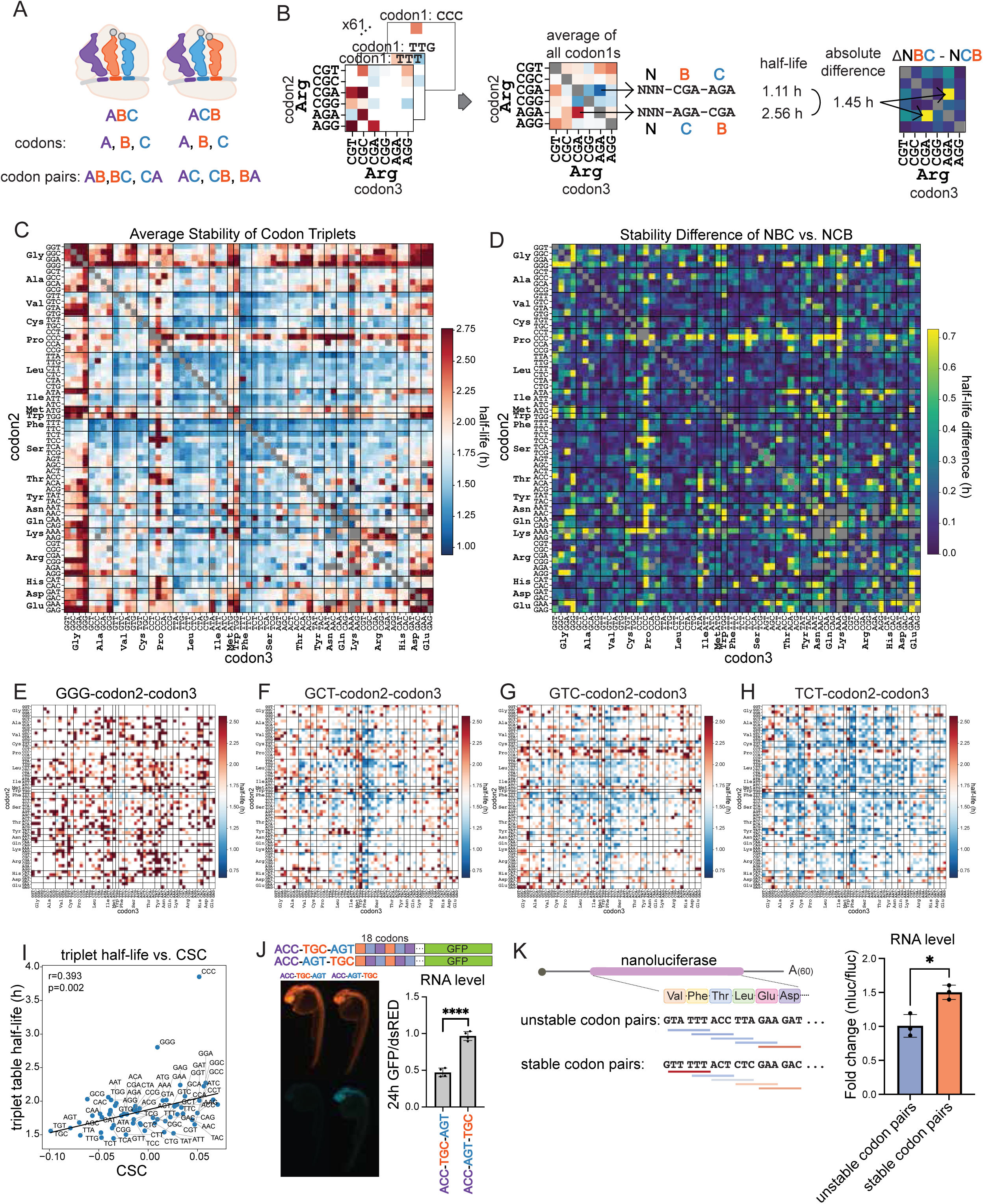
Codon pair stability depends on the codon order. **(A)** Schematics comparing reporters that have the same codon compositions but reversed codon pair orders. **(B)** Schematic of analysis: half-lives are averaged across all codon1s for each codon2-codon3, then the half-life difference is calculated between codon2-codon3 and codon3-codon2 (red, stable; blue, unstable; yellow, big half-life difference; purple, small half-life difference). **(C)** Heatmap of codon triplet half-lives, averaged across all codon1s and shown for each codon2-codon3 pair (red, stable; blue, unstable; gray, no data). See also Movie S1. **(D)** Heatmap representing the half-life difference between codon2-codon3 and codon3-codon2 (yellow, big half-life difference; purple, small half-life difference). Note that 1,588 out of 3,566 codon pairs have significantly different half-lives (q<0.05) from their inverted counterparts when we performed an inverse-variance-weighted two-sample t-test with a Benjamini-Hochberg correction. **(E-H)** Heatmap representing the half-life of codon1, codon2, and codon3, where codon 1 is fixed to GGG (E), GCT (F), GTC (G), or TCT (H) (red, stable; blue, unstable). **(I)** Scatter plot comparing average codon triplet half-life per codon to CSC (Spearman’s R). **(J)** Schematics comparing reporter constructs of codon triplets in ABC to ACB repeats, followed by GFP (top). Zebrafish embryo injected with codon triplet GFP and dsRED as a control. Fluorescence image at 24 hpf (bottom left) and RNA level quantification using qRT-PCR at 24 hpf (bottom right). Error bar represents standard deviation, and we performed an unpaired t-test: ****p<0.0001. **(K)** Schematics showing constructs encoding the same nanoluciferase with different codon pairs, based on the average half-life of codon triplets matrix (left). RNA level quantification using qRT-PCR at 6 hpf (right). Error bar represents standard deviation, and we performed an unpaired t-test: *p<0.05.

### Codon pair stability is dependent on translation

The reading frame specificity of codon pair stability (Fig. 1F) suggests that the ribosome, not the mRNA sequence per se, is the primary sensor of codon-pair context. We therefore hypothesized that removing translation initiation should eliminate codon-mediated mRNA stability. To test this, we generated matched reporter libraries in which the start codon was replaced with stop codons in all three frames (referred to as –start), to eliminate canonical translation initiation while preserving the coding sequence (Fig. 4A). Consistent with this hypothesis, –start reporters were significantly less stable than +start reporters (median: 0.86 h vs. 1.73 h, respectively; Fig. 4D–E) and their half-lives no longer correlated with CSC or tRNA levels, demonstrating that translation stabilizes mRNAs and that codon identity shapes stability only in the context of active translation (Fig. 4B–C).

**Figure 4.**
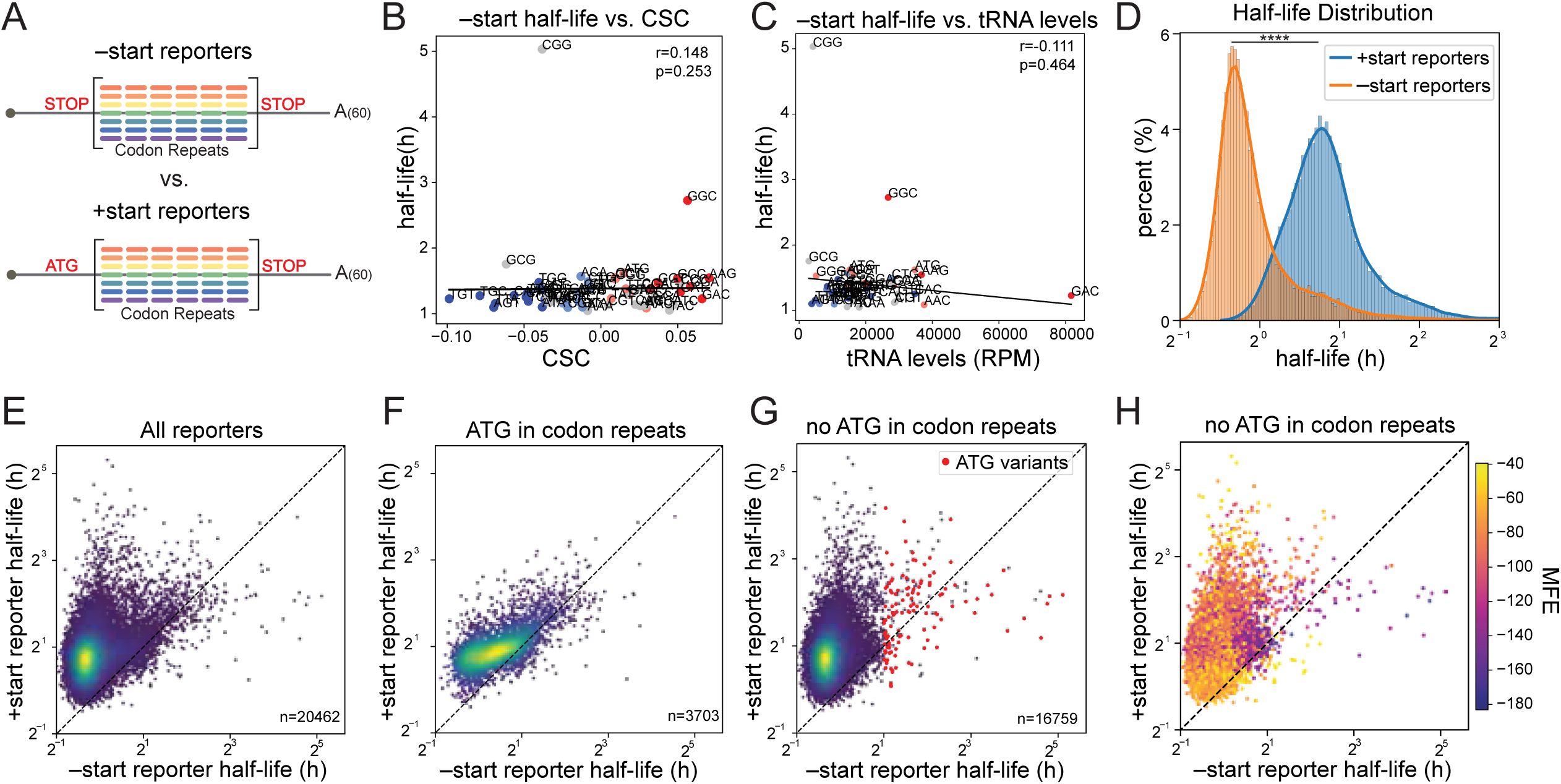
Codon pair stability is dependent on translation. **(A)** Schematics comparing massively parallel reporters that have the main start codon (+start reporters) to the ones that have the main start codon replaced with stop codons in all reading frames (–start reporters). **(B and C)** Scatter plot comparing –start reporters’ half-life per codon to CSC (Codon Stability Coefficient) (B) and tRNA levels in 5 hpf zebrafish embryo (C) (Spearman’s R). **(D)** Histogram of +start and –start reporters’ half-life (blue, +start reporters; orange, – start reporters). We performed a one-sided Wilcoxon signed-rank test: ****p<0.0001. **(E-H)** Scatter plots comparing half-lives of +start reporters to –start reporters of all reporters (E, n=20,462), reporters containing internal ATG in the CDS (F, n=3,703), reporters that do not contain internal ATG in the CDS (G, n=16,759; red, reporters contain one nucleotide variance from ATG and have half-life higher than 2h in –start reporters, n=128) (H; yellow, high minimum free energy (MFE); purple, low MFE). Note that reporters without internal ATG have a decreased half-life due to loss of translation at the main start codon. See also Figure S3.

Closer examination of the outliers revealed two groups of mRNAs that were stable despite loss of the canonical start codon. One group of reporters was enriched for internal ATGs and near-cognate initiators, indicating that downstream translation initiation preserved mRNA stability (Fig. 4F–G; Fig. S3D-I). A second group of reporters stabilized upon loss of translation and was predicted to be highly structured, as indicated by low minimum free energy (MFE). This suggests that ribosomes likely unwind these structures during elongation, exposing the RNA to decay^28,29^; when translation is blocked, the structure is preserved, thereby increasing mRNA stability (Fig. 4H). Together, these results demonstrate that CDS-mediated mRNA stability is translation-dependent, and reveal that RNA secondary structure encodes an orthogonal, translation-independent stability signal within the same sequence.

### Amino acids and amino acid pairs regulate mRNA stability through physicochemical and positional effects

To investigate whether amino acid identity, not just codon identity, shapes mRNA stability, we averaged reporter half-lives across synonymous codons for each amino acid (Fig. 5A). We found mRNA stability followed amino acid physicochemical properties: while amino acid size and charge showed a small trend but no significant correlation with mRNA half-life, mRNAs encoding hydrophobic amino acids were significantly less stable (Fig. 5B-D), suggesting that encoded amino acids can shape mRNA stability beyond individual codons.

**Figure 5.**
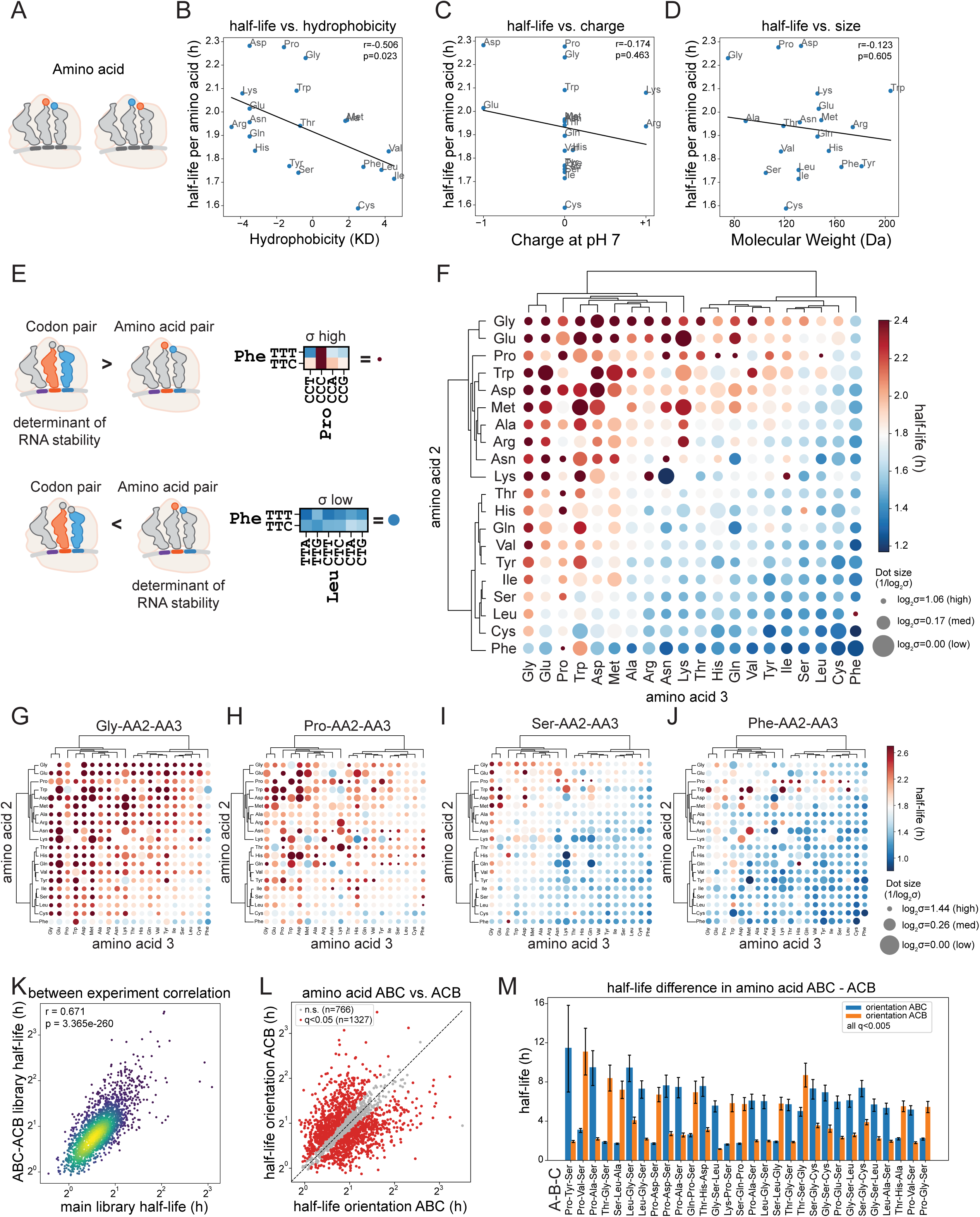
Amino acids and amino acid pairs regulate mRNA stability. **(A)** Schematics of amino acids in the translating ribosome showing how amino acids could influence mRNA stability. **(B-D)** Scatter plot comparing reporter half-life per amino acid to hydrophobicity (Kyte-Doolittle scale) (B), charge at pH 7 (C), and molecular weight (Da) (D) (Spearman’s R). **(E)** Schematic determining whether codon pairs or amino acid pairs drive mRNA stability by calculating the standard deviation of the codon pair half-lives coding for the same amino acid pair. **(F)** Heatmap representing average half-lives and standard deviations (on log2(half-life)) of the amino acid pairs calculated from codon triplets across all amino acid 1 (red, stable; blue, unstable; dot size, inverse of standard deviations). Note that half-lives and standard deviations of amino acid 2 - amino acid 3 compared to amino acid 3 - amino acid 2 are different. **(G-J)** Heatmap representing the half-life of amino acid1, amino acid2, and amino acid3, where amino acid1 is fixed to: Gly (G), Pro (H), Ser (I), Phe (J) (red, stable; blue, unstable; dot size, inverse of standard deviations). **(K)** Scatter plot comparing half-lives of the main codon repeat library and the subset ABC-ACB library (Pearson’s R). **(L)** Scatter plot comparing half-lives of the amino acid triplets in the orientation of ABC and ACB. We performed Welch’s t-test with Benjamini-Hochberg correction (gray, n.s.; red, p<0.05). **(M)** Top 30 half-life differences in amino acid triplets in the orientation of ABC and ACB. Error bars represent standard errors, and all 30 reporters show q<0.005 in Welch’s t-test with Benjamini-Hochberg correction.

To examine the influence of amino acid pairs on mRNA stability we calculated the average half-life and standard deviation across synonymous codon pairs encoding the same amino acid pair (Fig. 5E). We hypothesized that if codon pairs are the primary driver of mRNA stability, half-lives should vary widely among synonymous codon pairs encoding the same amino acid pair; conversely, if the amino acid pair drives stability, half-lives should remain consistent regardless of codon pair usage (Fig. 5E). For example, the average half-life for all synonymous codons encoding the phenylalanine-proline pair is high (half-life 3.59h), yet its constituent codon pairs span a wide range of half-lives (1.91-13.81h), indicating that in this case codon pair identity, rather than amino acid pair identity, is the dominant determinant mRNA stability (Fig. 5E). In contrast, phenylalanine-leucine consistently destabilizes mRNA (1.37h), with synonymous codon pairs being tightly clustered (1.20-1.61h), consistent with the amino acid pair driving this effect (Fig. 5E).

Reanalyzing average codon triplet stability by amino acid pair (Fig. 3C) resolved two clusters: hydrophilic amino acids stabilized mRNA, while hydrophobic amino acids destabilized it, consistent with the effect of individual amino acids (Fig. 5F). Indeed, holding constant the first amino acid and examining each pair revealed overall stabilizing to destabilizing effects, indicating that the first amino acid residue shifts the overall stability of the downstream pair (Fig. 5G-J).

To investigate the effect of the amino acid order on stability, we compared the half-lives of NBC versus NCB. Reversing amino acid pair order changed both the half-life and whether stability is primarily driven by the codons or the amino acids (Fig. 5F). For example, lysine-asparagine destabilizes mRNA with similar half-lives across synonymous codon pairs, suggesting that the destabilizing effect is driven by the amino acid pair, likely through peptide bond formation. Conversely, asparagine-lysine stabilizes mRNA, but with half-lives that vary widely across synonymous codons, indicating that this stabilizing effect is likely governed by the codon pair identity rather than amino acid sequence (Fig. 5F). To further probe the effect of the order of amino acids on mRNA stability, we designed a smaller targeted library of amino acid triplet repeats in two orientations: aa1-aa2-aa3 (ABC) and aa1-aa3-aa2 (ACB). mRNA half-lives correlated strongly with the main library, confirming reproducibility across experiments (Fig. 5K). Of 2,093 tested comparisons, 1,327 (63%) differed significantly in half-life, with fold changes up to 5.92 (Fig. 5L-M), indicating that the directional arrangement of amino acids shapes mRNA stability. Together, these results demonstrate that amino acid identity, pair composition, and order shape mRNA stability, revealing that the peptide sequence encodes a stability code beyond what is captured at the codon level.

### Machine learning identifies dominant determinants of mRNA stability

Having established that codons, codon pairs, amino acids, and amino acid pairs influence RNA half-life, we used machine learning to quantify their relative contributions. We trained models on the frequencies of these sequence features, together with MFE and GC/GC3 content, to predict reporter half-life (Fig. 6A). Sequence features predicted half-life well, with the gradient boosting explaining 73% of the variance and exceeding linear models (64%), indicating a contribution from nonlinear interactions between the different features (Fig. 6B, Fig. S4A). SHAP analysis of the gradient-boosted model ^30^ identified top predictors spanning all feature groups, revealing both stabilizing and destabilizing contributors (Fig. 6C). Several codons, codon pairs, amino acids, and amino acid pairs ranked among the top predictors (Fig. 6C), alongside the structural features MFE and GC/GC3 content (Fig. S4B-D).

**Figure 6.**
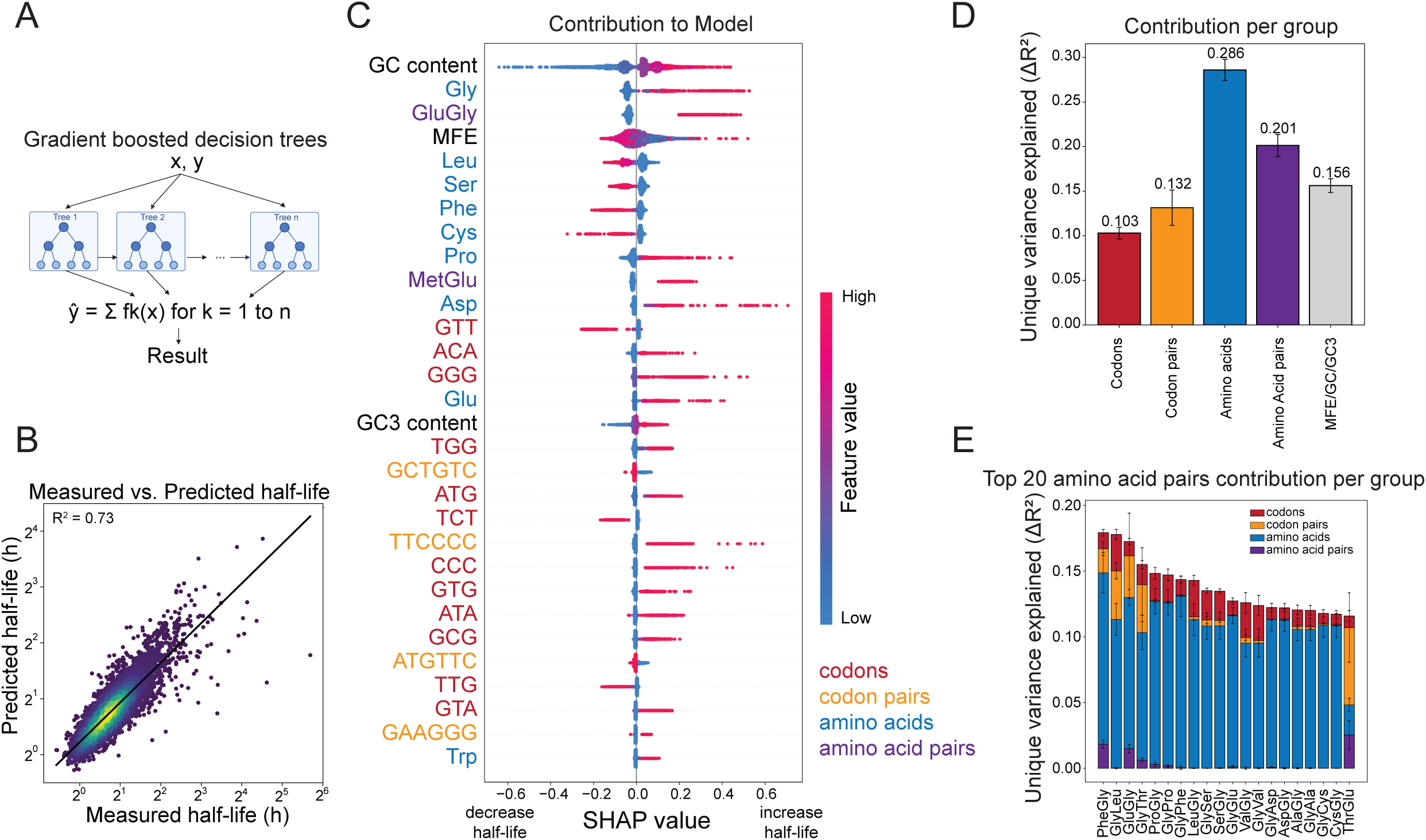
Machine learning identifies dominant determinants of mRNA stability. **(A)** Schematics illustrating machine learning model gradient boosted decision trees. **(B)** Scatter plot comparing the measured and predicted half-life of codon repeat MPRA of the model. **(C)** SHAP (SHapley Additive exPlanations) summary plot showing feature contributions to predicted mRNA half-life (Dot, reporters; positive SHAP, stabilizing; negative SHAP, destabilizing; pink, high feature value; blue, low feature value). **(D)** Variance partitioning by each feature group. The unique variance (ΔR^2^) of RidgeCV models fit on each feature set: codons (R^2^ codons - R^2^ amino acids), codon pairs (R^2^ codon pairs - R^2^ amino acid pairs), amino acids (R^2^ amino acids), amino acid pairs (R^2^ amino acid pairs - R^2^ amino acids), and GC/GC3 content/MFE. Error bars represent 95% CI across the 5 cross-validation folds. See also Fig. S4. **(E)** Per amino acid pair variance partitioning by each feature group. The top 20 contributing amino acid pairs are visualized. The unique variance (ΔR^2^) of RidgeCV models fit on each feature set: codons (R^2^ codons - R^2^ amino acids), codon pairs (R^2^ codon pairs - R^2^ amino acid pairs), amino acids (R^2^ amino acids), amino acid pairs (R^2^ amino acid pairs - R^2^ amino acids) (red, codons; orange, codon pairs; blue, amino acids; purple, amino acid pairs).

To determine how much of the variance in mRNA half-life each feature group explains, we applied nested linear decomposition, which accounts for the inherent dependencies among groups (e.g., codons encode amino acids, codon pairs encode amino acid pairs) to isolate each group’s unique contribution (Fig. 6D). Amino acid composition explained the greatest proportion of variance in mRNA half-life, followed by amino acid pairs, indicating that 49% of the mRNA stability was determined by the encoded peptide sequence (Fig. 6D). However, because synonymous codons encoding the same amino acids may underlie part of this signal, the contribution of amino acid identity and codon identity cannot be fully disambiguated; the 49% should therefore be interpreted as an upper bound on amino-acid-driven effects. Codon and codon pair usage contributed an additional 23% of variance independently of amino acid content (Fig. 6D). Partitioning variance for individual amino acid pairs showed that the contributions varied across amino acid pairs, indicating that stability is not governed by a single dominant code but emerges from the combined, context-dependent contributions of codon and amino acid identity (Fig. 6E). Together, these results show that mRNA stability is governed by multiple layers of coding sequence features, with both codon and amino acid pair context making independent and substantial contributions.

### Codon pairs regulate the stability of endogenous mRNAs

To assess the physiological relevance of codon pair stability, we examined whether codon pair composition is associated with endogenous transcript fate during development. During the maternal-to-zygotic transition (MZT), maternal mRNAs enriched in non-optimal codons are preferentially degraded ^9,10^. Based on these observations, we hypothesized that codon pair composition might also regulate maternal mRNA clearance. To test this hypothesis, we compared the log2 fold change of maternal transcripts across the MZT (log_2_(5.5h/0.75h)) ^31^. Transcripts with *miR-430* target sites, a key microRNA in maternal mRNA clearance ^5^, were the least stable, as expected. Remarkably, transcripts enriched in unstable codon pairs (bottom 25%) were degraded similarly to *miR-430* targets, while those enriched in stable codon pairs (top 25%) were more stable than either group (Fig. 7A). These results suggest that codon pairs regulate not only reporter stability but also endogenous mRNAs, playing a crucial role in maternal mRNA clearance during the MZT.

**Figure 7.**
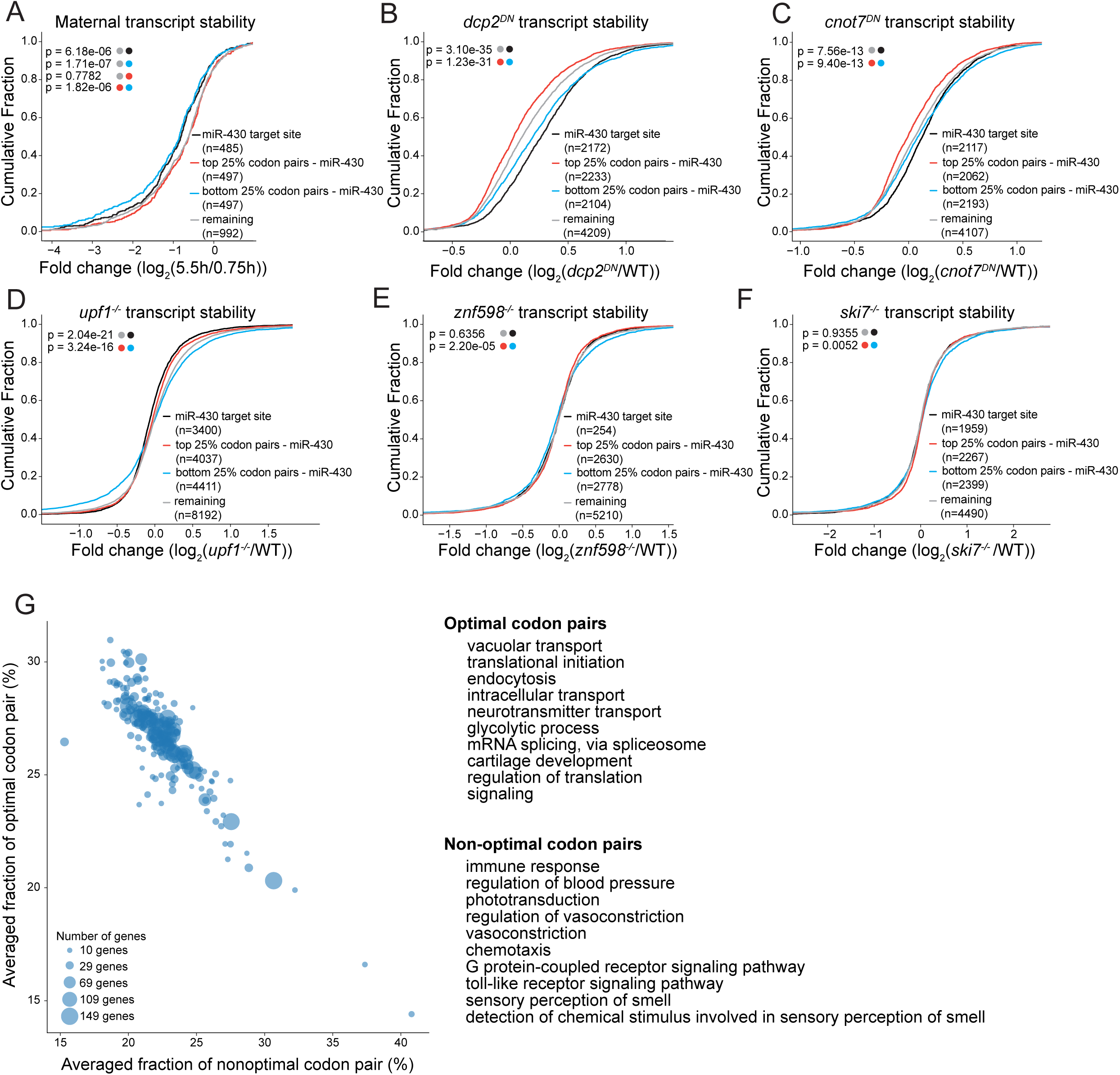
Codon-pair stability regulates developmental and biological functions via core cytoplasmic mRNA decay factors. **(A)** Cumulative distribution of mRNA abundance change (log_2_(5.5 hpf / 0.75 hpf)) for maternal genes classified by average codon-pair half-life and miR-430 target status. (black, transcripts with miR-430 target sites; red, transcripts with top 25% average codon pair half-life without miR-430 target sites; blue, transcripts with bottom 25% average codon pair half-life without miR-430 target sites; gray, remaining transcripts). Each p-value corresponds to the Kolmogorov-Smirnov test between the two distributions indicated by the colored dots. **(B-F)** Cumulative distributions of log2 fold change (dominant negative or knockout / wildtype) for genes in zebrafish embryos depleted of individual mRNA decay factors: Dcp2 (B), Cnot7 (C), Upf1 (D), Znf598 (E), and Ski7 (F) (black, transcripts with miR-430 target sites; red, transcripts with top 25% average codon pair half-life without miR-430 target sites; blue, transcripts with bottom 25% average codon pair half-life without miR-430 target sites; gray, remaining transcripts). Each p-value corresponds to the Kolmogorov-Smirnov test between the two distributions indicated by the colored dots. Note that Dcp2 and Cnot7 degrade unstable codon pair transcripts more than stable codon pair transcripts, while Znf598 and Ski7 do not. For Upf1, the difference depends on ORF length. See also Fig. S5. **(G)** Scatter plot comparing the mean fraction of optimal codon pairs and the mean fraction of nonoptimal codon pairs for each GO Biological Process term, using the codon-pair half-life (left). Each dot represents one GO term, and dot size indicates the number of annotated genes. The 10 GO terms most enriched for optimal codon pairs and the 10 terms most enriched for nonoptimal codon pairs are shown (right).

Having established that codon pairs regulate endogenous mRNA stability, we next sought to identify the decay machinery underlying this regulation. We reasoned that if a given factor mediates codon pair dependent decay, its knockout should preferentially stabilize transcripts enriched in unstable codon pairs compared to those enriched in stable codon pairs. To this end, we analyzed published RNA-seq data from loss-of-function studies of candidate decay factors, including the decapping enzyme Dcp2 ^32^, the deadenylase Cnot7 ^10^, the RNA surveillance factor Upf1 ^33^ involved in nonsense-mediated decay (NMD) and ORF-mediated decay (OMD; ^11^), and Znf598 ^34^ and Ski7 ^35^, both involved in no-go decay (NGD) (Fig. 7B-F). Dominant-negative Cnot7 preferentially stabilized transcripts enriched in non-optimal codon pairs, consistent with deadenylation as a critical step in codon-mediated decay. As a positive control, *miR-430* targets, which are cleared through deadenylation ^5^, were also stabilized (Fig. 7B-C; Fig. S5). Likewise, dominant-negative Dcp2 stabilized non-optimal codon pair transcripts and *miR-430* target sites. This stabilization was not observed in the poly(A) selected mRNAs, placing decapping downstream of deadenylation (Fig. 7B-C; Fig. S5A-F). Upf1 knockout showed a distinct pattern where long open reading frames enriched in non-optimal codon pairs were stabilized, indicating that poor translation of long non-optimal codon pair transcripts is surveilled by OMD (Fig. 7D; Fig. SG-J). In contrast, the no-go decay factors Znf598 and Ski7 did not preferentially stabilize unstable codon pair transcripts (Fig. 7E-F), indicating that NGD is not required for codon pair dependent decay. Together, these results indicate that codon pairs regulate endogenous transcript stability through deadenylation and subsequent decapping, with Upf1.

Finally, to assess whether codon pair optimality is associated with gene function, we calculated the fraction of optimal versus non-optimal codon pairs across the transcripts in each GO Biological Process term (Fig. 7G). Optimal codon pair genes were enriched for housekeeping functions such as translation, transport, and RNA processing, whereas non-optimal codon pair genes were enriched for inducible functions such as immune, vascular, sensory, and signaling pathways (Fig. 7G). These findings suggest that codon pair usage is tuned to gene function, conferring stability to constitutively expressed genes and rapid turnover to those requiring dynamic regulation.

## Discussion

Here, we show that codon pairs control mRNA stability beyond the effects of individual codons, in an order-dependent manner. The effect requires translation, implicating the ribosome as the reader of codon-pair context. Beyond codon pairs, the encoded amino acids and amino acid pairs add further context-dependent effects, and we quantified the relative contributions of each feature using machine learning. This regulation extends to the endogenous transcriptome, where codon pair usage shapes developmental gene regulation and biological function through deadenylation and subsequent decapping.

Our work reveals that the regulatory effect of a codon depends on its neighboring codons and identifies 58 synergistic and 165 antagonistic codon pairs relative to their individual codons, and that reversing codon pair order alters half-life in nearly half of all codon pairs (Fig. 2E, 3D). This set exceeds far beyond the 17 inhibitory codon pairs reported in yeast ^22^, suggesting that codon pair effects are more pervasive than previously appreciated. We further resolved a second layer of regulation. Amino acid pairs also shape mRNA stability, and by calculating the half-life spread of synonymous codon pairs encoding each amino acid pair (Fig. 5F), we estimated the relative contribution of codon pairs versus amino acid pairs, a step toward disentangling the regulatory codes embedded in the coding sequence.

What makes pair-level regulation order-specific and non-additive? The two classes of pairs we observe point to two distinct mechanisms. For pairs for which mRNA stability depends on synonymous codon pair choice, the stability difference likely reflects how the ribosome handles each codon-pair orientation at every decoding step, potentially through tRNA-tRNA interactions at adjacent ribosomal sites. Such tRNA-tRNA interactions could link mRNA stability to the differential isodecoder usage, which differs across developmental stages and tissues even when anticodon-level tRNA abundance remains unchanged ^27,36,37^. Pairs for which mRNA stability depends on the encoded amino acid pairs, the effect is better explained at the level of peptide bond formation or the nascent peptide. The rate of peptide bond formation depends on the identity and order of the P-site donor and A-site acceptor residues, and the growing nascent chain interacts with the ribosome exit tunnel in an order-dependent manner. The prominence of hydrophobicity among amino acid determinants (Fig. 5B) is consistent with nascent peptide interactions within the largely hydrophobic exit tunnel. In both cases, order specificity arises because reversing a pair places different tRNAs and amino acids in register at each step, consistent with the observed translation dependency (Fig. 4B-G). Quantitative measurements of elongation at specific codon pairs using approaches such as NaP-TRAP ^38^, will be needed to establish the kinetic basis of order-dependent stability.

How is codon pair context ultimately interpreted by the decay machinery? One explanation is that codon pair identity tunes the local rate of elongation, or the conformation of the ribosome, resulting in the recruitment of the deadenylation machinery ^13,16,39,40^. The model is supported by the factors required for degradation of the non-optimal codon pairs. Cnot7, a CCR4-NOT deadenylase component, and Dcp2, the decapping enzyme, were required to preferentially degrade endogenous transcripts with unstable codon pairs. Upf1 was additionally required for long open reading frames with non-optimal codon pairs, consistent with the ORF-mediated decay of poorly translated transcripts ^11^. In contrast, no-go decay factors Znf598 and Ski7 were not required (Fig. 7B-F). By coupling slow elongation to deadenylation and decapping, codon pair regulation of stability is likely mediated by the mechanism established for single-codon optimality. Future structural studies using cryo-EM to capture the relevant ribosome conformations and mass spectrometry will be needed to identify associated factors with codon pair-mediated regulation.

Our results indicate that codon pair-mediated mRNA stability shapes gene regulation. During early development, maternal mRNA clearance at the MZT is an indispensable step. miR-430 clears several hundred target maternal mRNAs ^5^, and the non-optimal codon usage in maternal transcripts further contributes to their decay ^9,10^. Our analysis shows that transcripts enriched in unstable codon pairs are degraded as efficiently as miR-430 targets, as measured by log2 fold change in transcript abundance across MZT, revealing a role for codon pairs, alongside single codons, in programmed maternal mRNA clearance (Fig. 7A). This link between codon pair usage and gene regulation expands across the transcriptome. Because the encoded peptide sequence accounts for a substantial fraction of the CDS-encoded stability signal, though part of this may reflect the codons encoding those amino acids (Fig. 6D), we propose that the function of the encoded protein influences a transcript’s baseline turnover rate over evolutionary selection, while codon and codon pair usage modulate it as an additional layer of fine-tuning. It will be interesting to extend this framework to other organisms and cell types, which will let us understand the codon pair stability code across different species.

Beyond its biological role, mRNA stability regulation at the level of codon pairs offers a practical design principle for mRNA therapeutics and vaccines, where transcript stability governs both protein output and durability. Current codon optimization strategies are largely based on single-codon optimality ^41^. Our results add order-specific codon pair identity as an additional design tool, one that can be tuned without altering the encoded protein (Fig. 3K). Codon pair design can stabilize mRNA for sustained expression or destabilize it for transient, tightly controlled expression or live-attenuated vaccine design, expanding the tunable range available for therapeutics. Together, our results reveal a new layer of gene regulation in which codon and amino acid order shape mRNA stability, with broad implications for gene regulation and RNA therapeutics.

### Limitations of the Study

Several limitations of this study should be acknowledged. Our MPRA reporters share identical UTRs to isolate CDS effects from confounding elements on stability, and thus do not capture potential interactions between the coding sequence and UTR regulatory elements^10,42,43^. Our short repeat-based coding sequences limit the nascent chain to the ribosome exit tunnel, resulting in a functional analysis focused on ribosome-driven clearance, yet the longer nascent peptides of endogenous transcripts may engage additional co-translational quality control mechanisms^44^. Finally, our measurements were made using in vitro transcribed mRNAs that lack RNA modifications with fixed poly(A) tail length and were tested in zebrafish embryos within a defined developmental window; while these reporters isolate CDS-driven effects, they may not fully recapitulate the behavior of endogenous transcripts in their native genomic and cellular context with endogenous RNA modifications and splicing. Future studies will be needed to determine the extent to which the codon pair stability code operates across other cell types, organisms, and, critically, in human cells, where its implications for gene regulation and therapeutic mRNA design are most directly relevant^18,45^.

## Acknowledgments

We thank D. Olowookere, C. Hickey, D. Karpel, S. Dube, T. Gerson for technical help; C. Castaldi, E. Sykes, B. Carlson, C. Goncalves, M. Singh, and B. De Kumar from the Yale Center for Genome Analysis for sequencing support; L. Weiss and C. Boswell for feedback on the manuscript; A. Bazzini and all members of the Giraldez laboratory for intellectual and technical support. This research was supported by the National Institute of Health (grants R01 HG013516, R01 HD119238, R35 GM122580, to AJG).

## Author Contributions

HL, JDB, MAM and AJG conceived the project. HL performed experiments and analysis with the support of DM and CEV. SK and ECS contributed experimental analysis and intellectual input. CMT generated dominant negative embryos and performed RNA-seq, and TDN performed RNA-seq and GO term analysis. HL and AJG wrote the manuscript.

## STAR Methods

## Key Resources Table

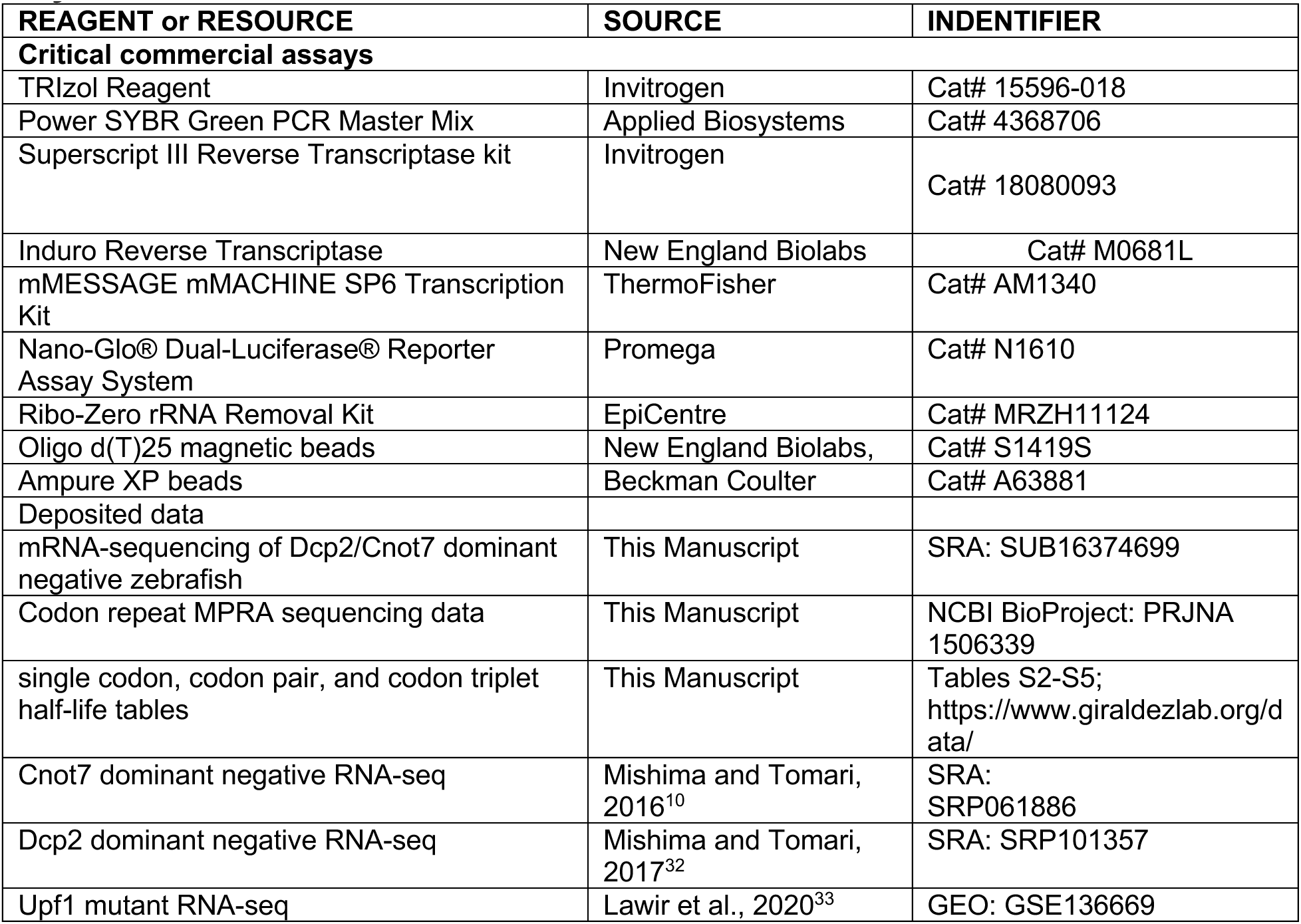
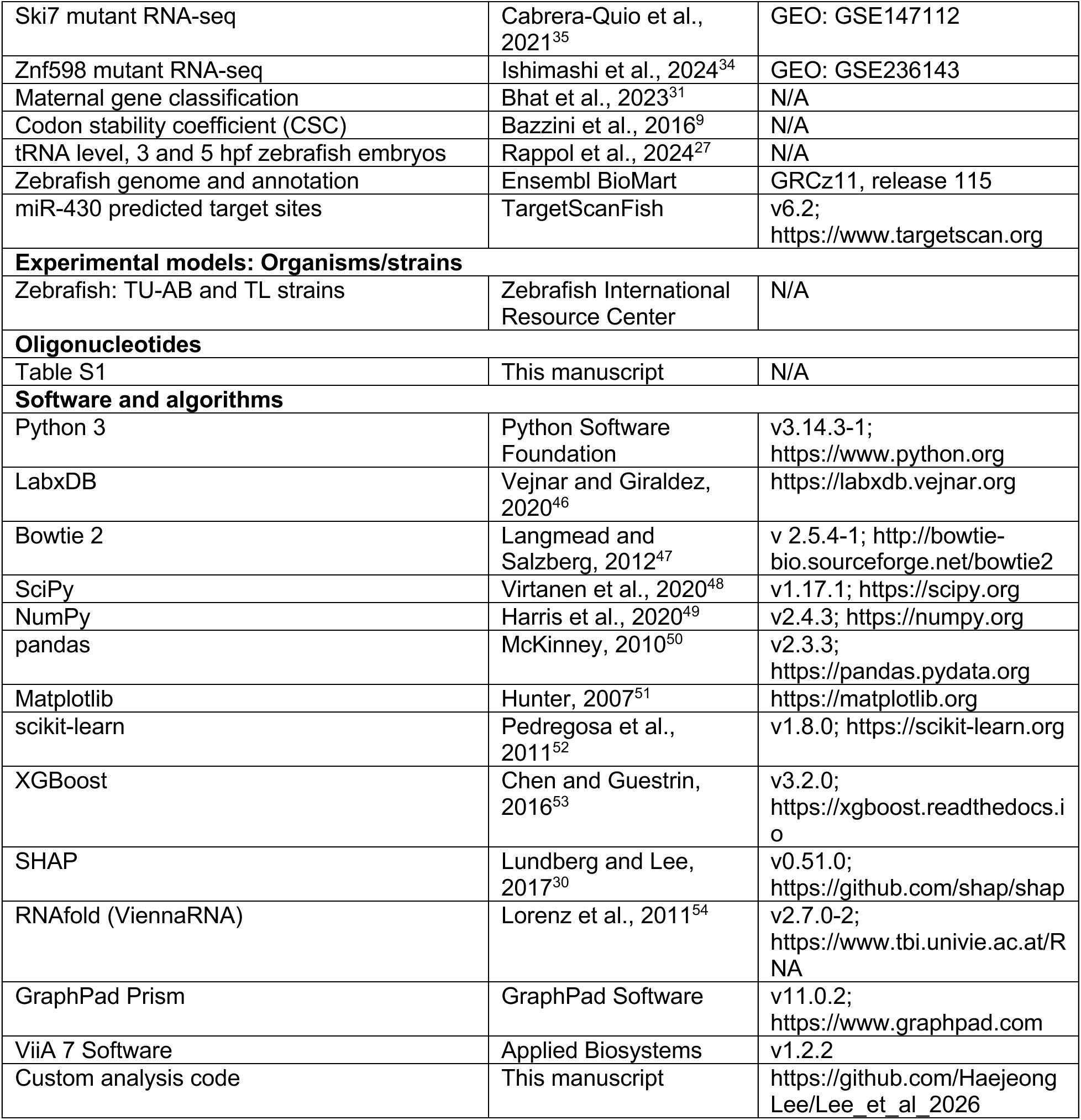

## RESOURCE AVAILABILITY

## CONTACT FOR REAGENT AND RESOURCE SHARING

Further information and requests for resources and reagents should be directed to and will be fulfilled by the Lead Contact, Antonio J. Giraldez

### Data availability

The data supporting the findings of this study are available from the corresponding authors upon request. The sequencing data generated in this study have been deposited in NCBI under the BioProject ID PRJNA1506339. The processed half-life tables from the sequencing data are available in Supplementary Data 2-4 or on the Giraldez Lab datahub (https://www.giraldezlab.org/data/).

### Code availability

All scripts used to process and analyze data will be released at the time of publication (https://github.com/HaejeongLee/Lee_et_al_2026).

## EXPERIMENTAL MODEL AND STUDY PARTICIPANT DETAILS

### Zebrafish embryo production

Zebrafish wild-type embryos were obtained from natural matings of 12-month-old adult zebrafish of mixed wild-type backgrounds (TU-AB and TL strains). Wild-type adults were selected randomly for mating. Zebrafish were maintained in accordance with AAALAC research guidelines, under a protocol approved by Yale University IACUC. All zebrafish and embryo experiments were carried out at 28°C. Embryos were grown and staged according to standard protocols to ensure all embryos were at the same expected developmental stages before sample collection. Embryos at developmental stages between 3 and 24 hours post-fertilization (hpf) were randomly collected per sample, as specified across different experiments.

## METHOD DETAILS

### Massively parallel reporter design and *in vitro* transcription

Codon reporters were generated as follows: synthetic oligos encoding 42 codon stretches of single codons, codon pairs, codon triplets, and codon combinations, each carrying common and distinct barcodes, were ordered from GenScript and IDT (Integrated DNA Technologies). Using overlap PCR extension, SP6 and Illumina 5’ adaptor sequences were inserted in the 5’ end of the synthetic oligo, and beta-globin 3’ UTR followed by 60A were inserted in the 3’ end of the synthetic oligo. Final constructs were size-purified on a 2% agarose gel and used as templates for in vitro transcription with the SP6 mMessage mMachine kit (ThermoFisher, AM1340) to generate reporter mRNA. Transcribed mRNA was purified using the Monarch RNA Cleanup kit (New England Biolabs, T2040L).

Four MPRAs were used in this study: (1) the main codon repeat libraries, comprising single codon, codon pair, and codon triplet repeats, (2) targeted single codon repeat and codon pair libraries, injected and sequenced separately to increase UMI depth, (3) matched –start libraries, in which the main start codon was replaced with stop codons in all three reading frames (TAGGTAGCTAG), and (4) targeted amino acid triplet libraries, in which three amino acids were encoded in ABC and ACB orientations.

### Embryo injections

All injections and drug treatments were carried out on wild-type one-cell stage dechorionated embryos. Varying amounts of reporter mRNA per embryo and collections were used as follows: (1) 12.5pg of codon repeat MPRA mRNA were injected, 25 embryos per replicate were collected at 3, 4, 5, 6, 7, and 8 hpf, (2) 0.5 pg of nanoluciferase mRNA with 19.5 pg of firefly luciferase mRNA were injected, 5 embryos per replicate were collected at 6hpf, (3) 50pg of 6x codon triplets-GFP mRNA with 50pg of dsRED mRNA were injected, 5 embryos per replicate were collected at 24hpf, and (4) 75pg of Dcp2 and Cnot7 dominant negative mRNA ^55^ were injected, 25 embryos per replicate were collected at 6hpf.

### Sequencing library preparation

Total RNA was extracted using Trizol (Invitrogen, 15596-018) from the collected samples, five spike-ins designed based on the External RNA Controls Consortium (ERCC) were added. For the Dcp2Cnot7 dominant negative RNA-seq, rRNA removal using Ribo-Zero rRNA Removal Kit (EpiCentre, MRZH11124) or poly(A) selection using Oligo d(T)25 magnetic beads (New England Biolabs, S1419S) was performed. Then RNA was reverse transcribed using Superscript III kit (Invitrogen, 18080085) or Induro Reverse Transcriptase (New England Biolabs, M0681L). cDNA was size-purified using AMPure XP beads (Beckman Coulter, A63881) and purified again after amplifying the inserted codon repeats directly using Illumina 5’ and 3’ adaptors with UMI that bind to the reverse transcription primer. Libraries were sequenced on Illumina HiSeq 2000, Illumina NovaSeq 6000, and Illumina NovaSeq X Plus with paired-end 150bp. Oligonucleotides used for RT and PCR are listed in Table S1.

### Sequencing analysis and half-life calculation

Sequencing data were stored using LabxDB ^46^. Paired-end reads were trimmed and demultiplexed using ReadKnead (https://github.com/vejnar/ReadKnead). Barcodes identifying replicates and UMIs were extracted from read 2. Trimmed reads were mapped using Bowtie2 ^47^. To count only perfect-match reads of highly repetitive sequences, insert sequences were reconstructed from reads 1 and 2 and were compared with the design sequences. UMI counts of constructs were normalized using five spike-ins. Only reporters with UMI counts at more than three time points between 3 and 8 hpf across three biological replicates, and with a total UMI count across all time points greater than 400, were used in the half-life calculation. mRNA abundance over time was modeled as first-order exponential decay:

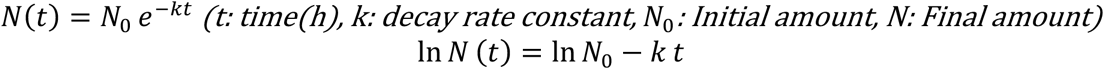

For each reporter, half-life was computed from the decay rate, and the standard error of half-life was calculated by first-order error propagation:

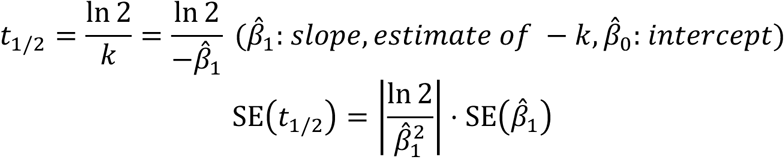

After calculating half-lives, we discarded 3% of reporters whose linear regression fitting was poor based on the standard error of the slope. No other data were excluded from the analyses.

### Calculation of sequence and physicochemical features

Codon stability coefficients (CSC) were taken from Bazzini et al. (2016)^9^. tRNA abundances at 3and 5hpf were taken from Rappol et al. (2024)^27^ and expressed as reads per million (RPM). Minimum free energy (MFE) was computed for each reporter sequence using RNAfold from ViennaRNA v2.7.0-2^54^, with default parameters at 28°C. GC and GC3 contents were calculated as the fraction of G or C across all position and across third codon positions, respectively. Amino acid hydrophobicity was taken from the Kyte-Doolittle scale ^56^, charge was assigned at pH 7 ^57^, and molecular weight was taken in daltons ^58^.

### Fluorescence microscopy imaging

Embryos injected with 6x codon triplet-GFP and dsRED mRNA were imaged at 24 hpf. Fluorescence images were acquired using a Zeiss SteREO Discovery.V8 stereomicroscope equipped with an Achromat S 1.0x objective and AxioCam MR R3 camera. GFP and DsRed fluorescence were imaged separately using the appropriate filter sets. Identical exposure times and acquisition settings were used for all embryos within the experiment.

### qRT-PCR measurements of RNA abundance

Total RNA was extracted from 5 embryos per experimental condition and DNase-treated. cDNA was synthesized from total RNA using reverse transcription with Oligo(dT)_12-18_ Primer (Thermo Fisher Scientific, 18418012) and the Superscript III reverse transcriptase kit (Invitrogen, 18080093). cDNA was diluted 1:20, and 10 μL PCR reactions were prepared with 5 μL of Power SYBR Green PCR Master Mix (Applied Biosystems, 4368706), 4.5 μL of diluted cDNA, and 0.5 μL of 10 μM forward and reverse primer mix. At least three biological and three technical replicates were performed for each sample. Relative expression was measured with ViiA 7 software v1.2.2 using the ΔΔCT method. Oligonucleotides used for qRT-PCR are listed in Table S1.

### Design of luciferase sequences based on codon pair stability

Given a target protein sequence of N amino acids, we designed the synonymous coding sequence in which adjacent codon pairs were either maximally stabilizing or maximally destabilizing. Average codon triplet half-lives as the scoring metric for dynamic programming to optimize codon pairs ^59^. Designed nanoluciferases were ordered (Twist Bioscience) with constant 5’ UTR and 3’ UTR fragments, then amplified using SP6 primers and 60A primers. Oligonucleotides used for assembly of nano luciferase are listed in Table S1.

### Machine learning and variance partitioning

Reporter half-lives were log2 transformed and used as the prediction target. Features comprised the within-reporter frequencies of the 61 codons, 3,721 codon pairs, 20 amino acids, and 400 amino acid pairs, together with MFE, GC content, and GC3 content.

Reporters were split into training and held-out test sets (80/20, random_state=42). Six models were compared on the identical split: Gradient Boosting, Random Forest, Ridge, Lasso, and Elastic Net. Learnings were performed using Python3 with scikit-learn 1.8.0, XGBoost 3.2.0, NumPy 2.4.3, SciPy 1.17.1, and pandas 2.3.3. Feature contributions to the gradient-boosted model were quantified using SHAP (SHapley Additive exPlanations) values ^30^, computed with shap 0.51.0.

To estimate the independent contribution of each feature group while accounting for nesting between groups (codons encode amino acids and codon pairs encode amino acid pairs), unique variance was partitioned using RidgeCV models fit on each feature set and assessed by regressing log2 (half-life) on the out-of-fold prediction. Unique variance was defined as ΔR² = R² (codons) - R² (amino acids) for codons, ΔR² = R² (codon pairs) - R² (amino acid pairs) for codon pairs, ΔR² = R² (amino acid pairs) - R² (amino acids) for amino acid pairs, and the model R² for amino acids and for the combined MFE/GC/GC3 set.

### Calculation of average codon pair half-life of endogenous transcripts

Zebrafish transcript sequences were obtained from Ensembl BioMart (GRCz11, release 115). Transcripts were retained only if the CDS began with an ATG start codon and ended with a canonical stop codon (TAA, TAG, or TGA). Transcripts containing ambiguous nucleotides were excluded. For genes with multiple isoforms, the transcript encoding the longest open reading frame was selected as the representative transcript. Codon-level mRNA half-life values were derived from the average of the codon triplet library. For each gene, codon-pair half-life values were averaged across all consecutive codon pairs in the CDS. We collected maternal genes from Bhat et al. (2023)^31^ and calculated log2 fold changes in expression between 5.5 hpf and 0.75 hpf. miR-430 target genes were identified using TargetScan (Fish 6.2, https://www.targetscan.org). For decay factor analysis, we used RNA-seq datasets from zebrafish embryos with individual decay factors perturbed: Dcp2 and Cnot7 ^10,32^, Upf1 ^33^, Ski7 ^35^, and Znf598 ^34^. For the ORF length analyses of Upf1 knockout RNA-seq, transcripts were ranked by CDS length and separated into top 25% and bottom 25%.

### GO analysis for codon optimality composition

For each zebrafish gene, individual codon pairs in the CDS were classified as optimal (top 25% of the half-life distribution) or nonoptimal (bottom 25% of the half-life distribution), using the codon-pair half-life values defined above. For each gene, the fraction of optimal codon pairs and the fraction of nonoptimal codon pairs in the CDS were calculated.

Genes with GO Biological Process terms were retrieved from Ensembl BioMart (GRCz11, release 115). For each GO term with at least 10 annotated genes, the mean fraction of optimal codon pairs and the mean fraction of nonoptimal codon pairs were calculated. GO terms were ranked by the difference between their mean optimal and nonoptimal fractions. The 10 terms with the largest positive difference (most enriched for optimal codon pairs) and the 10 terms with the largest negative difference (most enriched for nonoptimal codon pairs) were shown.

## QUANTIFICATION AND STATISTICAL ANALYSIS

### Statistical analyses

The plots and statistical analyses in Fig. 3J, K were generated using GraphPad Prism 11.0.2. All other analyses, unless otherwise stated, were performed using custom scripts written in Python 3. Plots were generated using the Matplotlib package ^51^. Statistical analyses were performed using the SciPy 1.17.1 ^48^ and NumPy 2.4.3 ^49^ packages. No statistical methods were used to predetermine sample size. Reporters with poor exponential decay fits (3% of the library) were excluded, as described above. No other data were excluded from the analyses. The experimental findings were verified by independent experimental replicates as indicated in figure legends and text. The experiments were not randomized, and investigators were not blinded to allocation during experiments and outcome assessment.

## Supplemental Video and Excel Table Title and Legends

**Movie S1.** *Half-lives of codon triplets across different codon 1.* Related to Figure 3. Shown are codon 2 - codon 3 half-life heatmaps across all 61 codon 1.

**Supplementary Table 1.** Oligonucleotide sequences utilized in the manuscript.

**Supplementary Table 2.** Half-life data of main (+start) codon repeat MPRA library. Related to Figure 1-7.

**Supplementary Table 3.** Half-life data of targeted single codon and codon pair repeat MPRA library. Related to Figure 1-2, 4, 6.

**Supplementary Table 4.** Half-life data of –start codon repeat MPRA library. Related to Figure 4.

**Supplementary Table 5.** Half-life data of targeted amino acid triplet. Related to Figure 5.

**Supplementary Table 6.** Average half-life and standard deviation across synonymous codon pairs encoding the same amino acid pair to distinguish determinants of mRNA stability. Related to Figure 5.

## Supplementary Figure Titles and Legends

**Figure S1.**
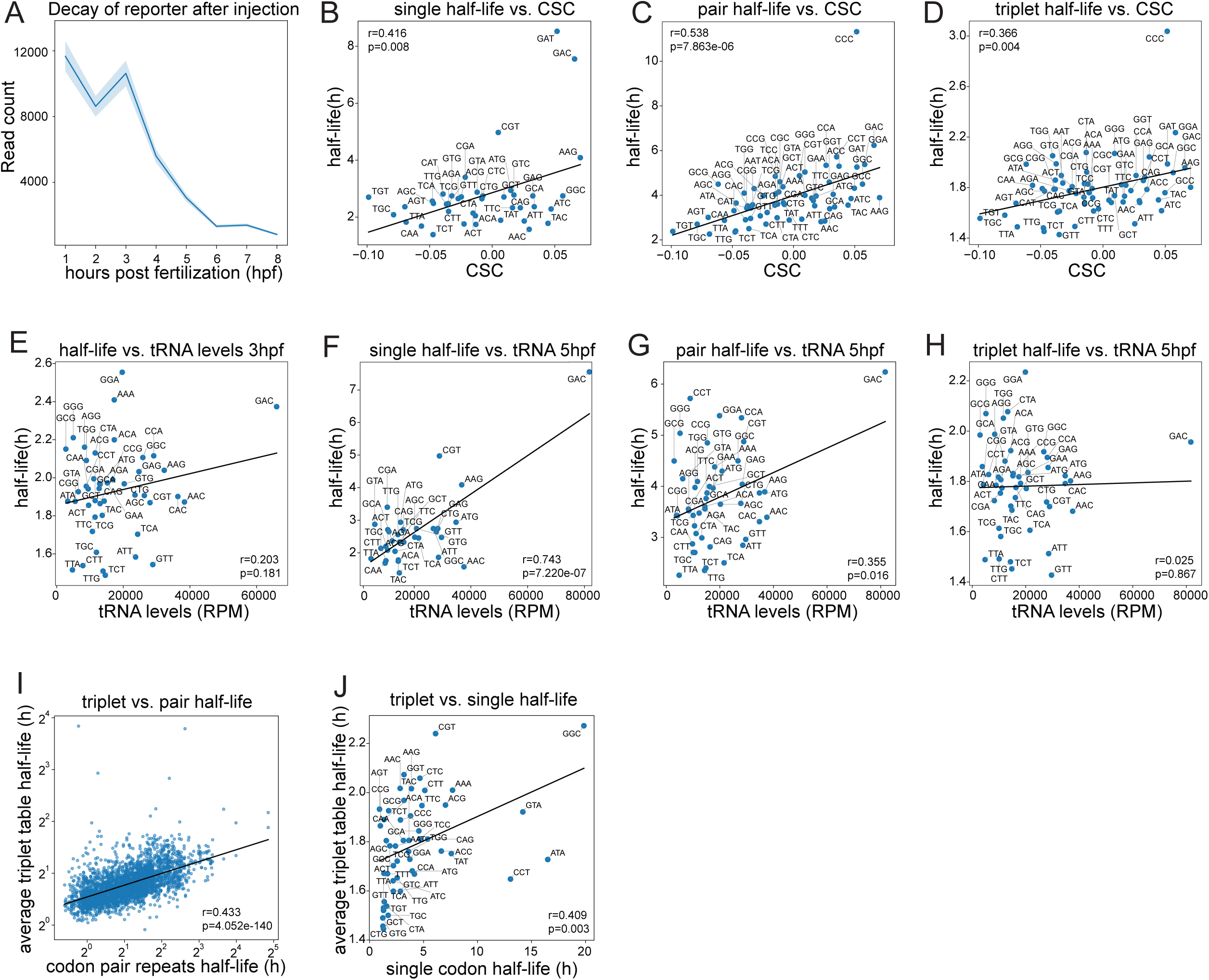
Codon repeat MPRA half-life correlates with CSC and tRNA levels. **(A)** Time-course decay of reporter read counts from 1 hpf to 8 hpf. Reporter read counts were stable until the onset of the MZT at around 3hpf, after which they decayed exponentially (Line, mean; shaded band, SEM). **(B-D)** Scatter plot comparing reporter’s half-life to CSC (Codon Stability Coefficient). Per-codon average of half-lives was calculated from the single (B), pair (C), and triplet (D) codon repeats (Spearman’s R). **(E-H)** Scatter plot comparing reporter’s half-life to tRNA levels, using all reporters against 3 hpf tRNA levels (E) and single (F), pair (G), and triplet (H) codon repeat reporters against 5 hpf tRNA levels (Spearman’s R). Note that the more complex the library becomes, the less correlation with tRNA level is observed, indicating extra layers of mRNA stability regulation steps in more complicated coding sequences. **(I-J)** Scatter plot comparing average half-lives of triplet codon repeat reporters and pair codon repeat reporters (I) and single codon repeat reporters (J) (Pearson’s R).

**Figure S2.**
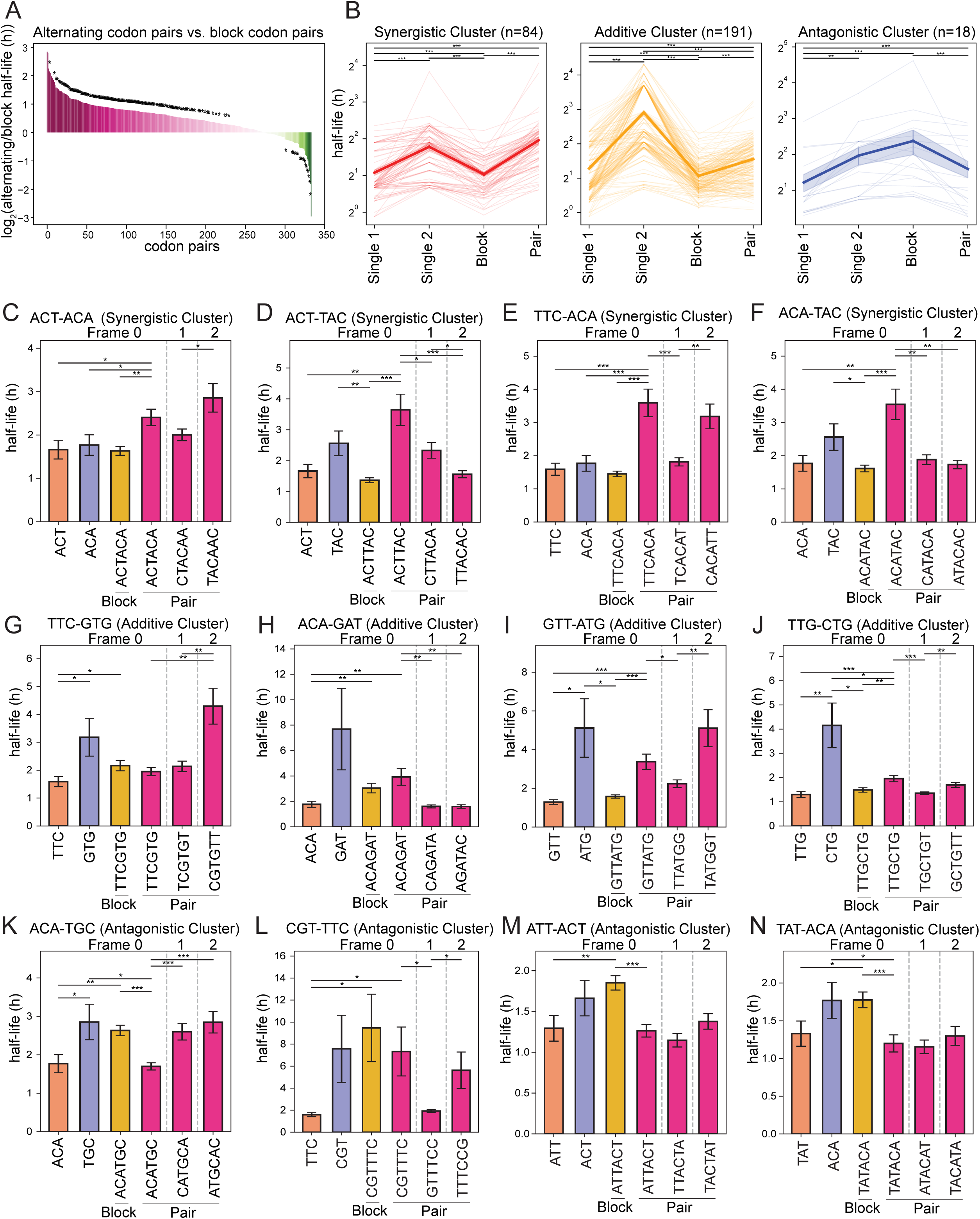
Codons arranged in pairs or blocks show different mRNA stability. **(A)** Half-life difference between alternating codon pair reporters and block codon reporters that have the same number of codons arranged differently (pink, synergistic; green, antagonistic). Note that 190 out of 334 codon pairs (57%) tested have significantly different half-lives. We performed Welch’s t-test: *p<0.05. **(B)** Clustering of half-life patterns across single-codon repeats (Single 1, Single 2), two-codon mixtures in block versus pair arrangements (thick line, mean; shaded band, SEM; thin line, individual reporter sets). We clustered reporter sets that have half-lives for both two-codon mixtures and at least one single codon. We performed k-means clustering with k=3 and Fisher’s combined Welch’s t-test: *p < 0.05, **p <0.01, ***p < 0.001. **(C-N)** Half-life comparison of reporters carrying single-codon repeats, two-codon mixtures in block versus pair arrangements showing synergistic (C-F), additive (G-J), and antagonistic (K-N) effects in alternating codon pairs compared to block codon pairs. We performed Welch’s t-test: * p < 0.05, **p < 0.01, ***p < 0.001.

**Figure S3.**
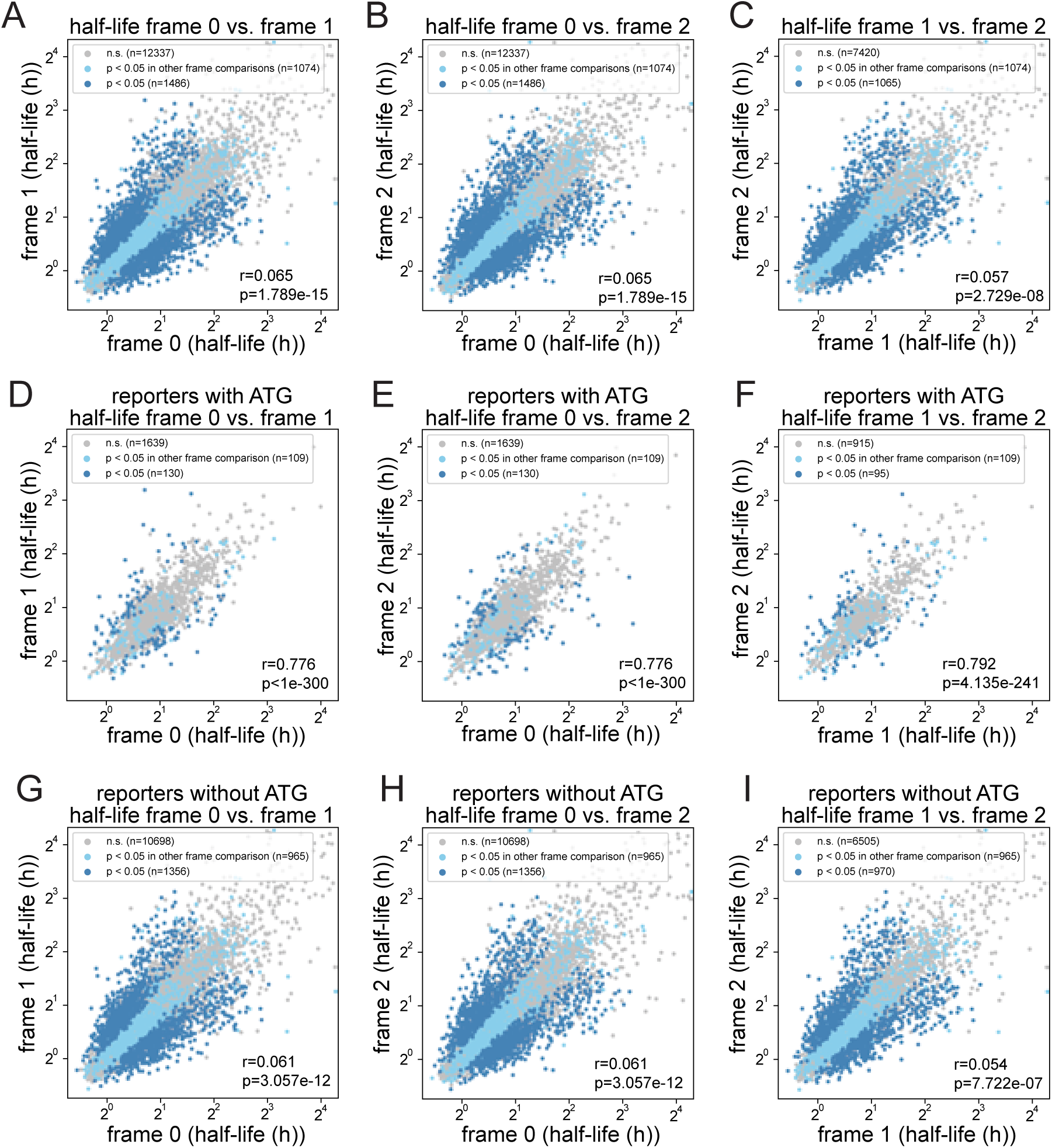
Codon repeat MPRA half-life is specific to the reading frame. (A-C) Scatter plot comparing half-lives of all codon repeat MPRA in the original reading frame and +1 reading frame (A), +2 reading frame (B), and +1 reading frame with +2 reading frame (C). Note that the correlation of reporter half-lives between different reading frames is near 0 (blue, Benjamini-Hochberg-corrected Welch’s t-test p < 0.05; light blue, Benjamini-Hochberg-corrected Welch’s t-test p < 0.05 in the other frame comparison; gray, not significant; Pearson’s R). **(D-F)** Scatter plot comparing half-lives of codon-repeat MPRA containing ATG in the coding sequence in the original reading frame and +1 reading frame (D), +2 reading frame (E), and +1 reading frame with +2 reading frame (F). Note that half-lives of reporters containing ATG have high correlations between different reading frames, possibly due to alternative translation initiation (blue, Benjamini-Hochberg corrected Welch’s t-test p < 0.05; light blue, Benjamini-Hochberg corrected Welch’s t-test p < 0.05 in other frame comparison; gray, not significant; Pearson’s R). **(G-I)** Scatter plot comparing half-lives of codon-repeat MPRA that do not contain ATG in the coding sequence in the original reading frame and +1 reading frame (G), +2 reading frame (H), and +1 reading frame with +2 reading frame (I). (blue, Benjamini-Hochberg corrected Welch’s t-test p < 0.05; light blue, Benjamini-Hochberg corrected Welch’s t-test p < 0.05 in other frame comparison; gray, not significant; Pearson’s R).

**Figure S4.**
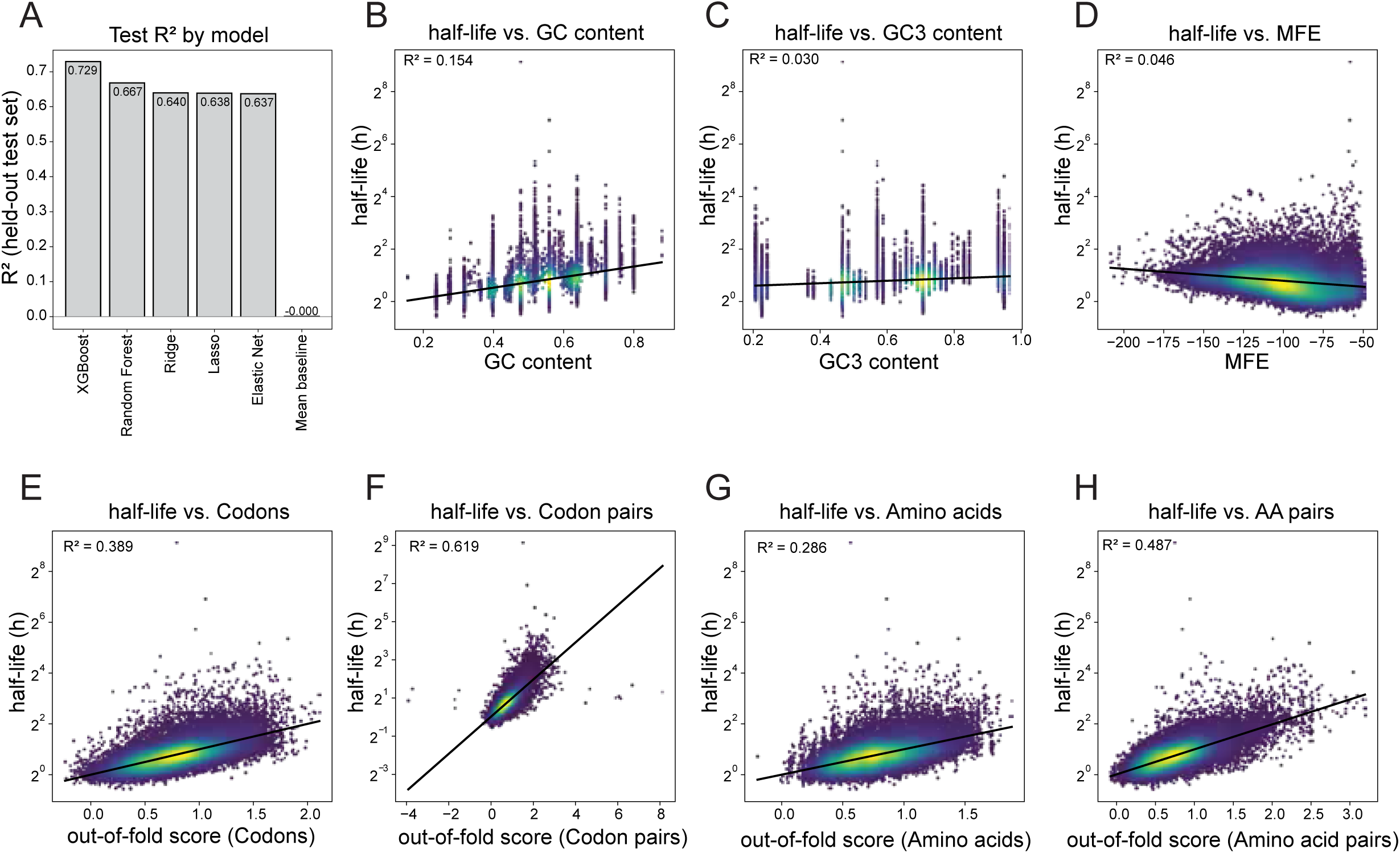
Each feature group is associated with reporter mRNA half-life. **(A)** Bar plot comparing test R^2^ score for each model. Every model is trained and tested on the same held-out split. The mean baseline is the prediction made using the mean of the training set. Note that gradient boosting was the best predictor among the models tested. **(B-H)** Scatter plot comparing reporter half-life and feature group GC content (B), GC3 content (C), MFE (D), out-of-fold score of codons (E), codon pairs (F), amino acids (G), and amino acid pairs (H). R^2^ represents the fraction of the variance in log2(half-life) explained. R^2^ values here are raw, non-nested associations for each feature group individually and do not account for the dependency between groups. Unique contributions after nesting are shown in Figure 6D.

**Figure S5.**
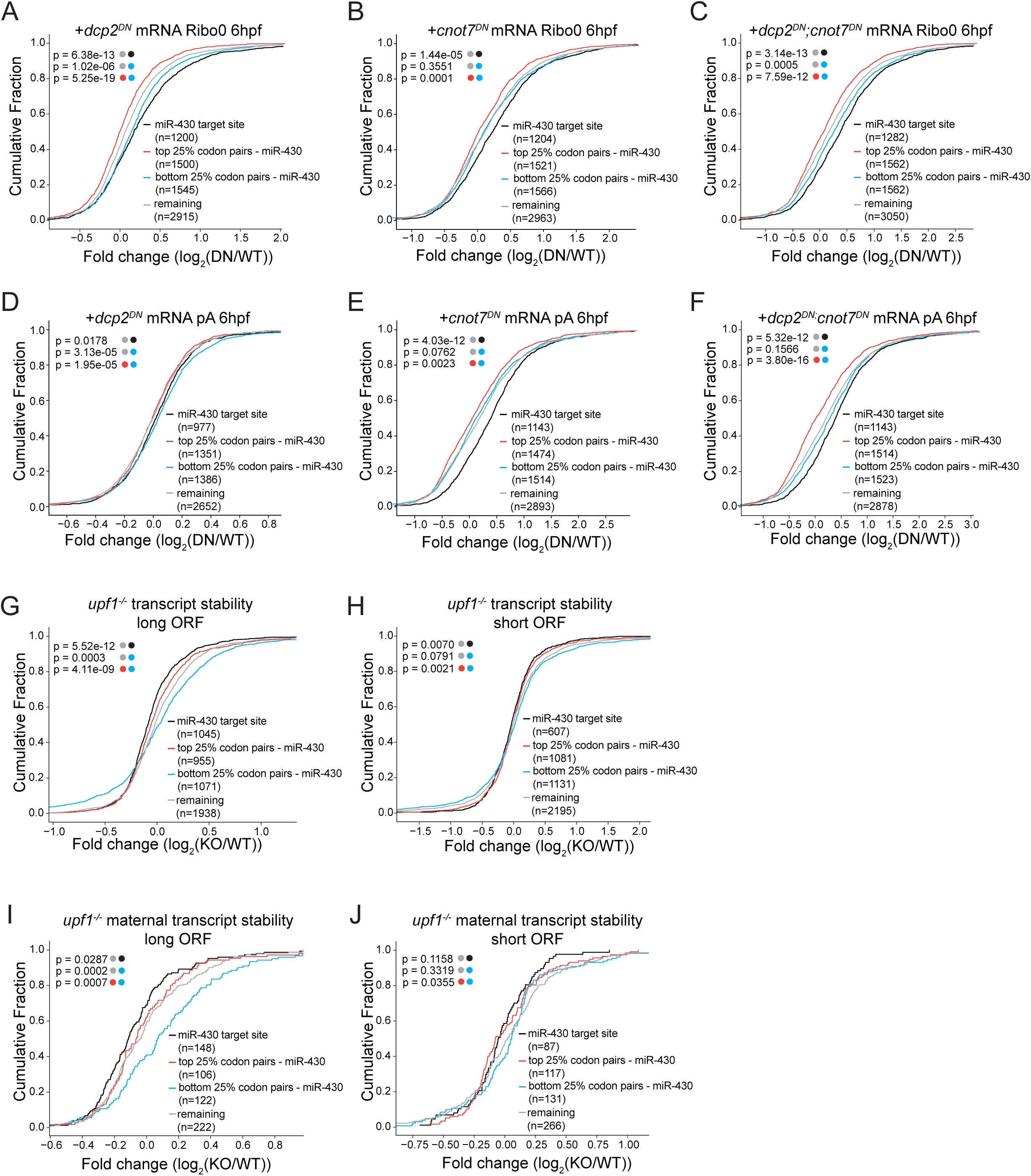
Dcp2, Cnot7, and Upf1 preferentially destabilize transcripts enriched in unstable codon pairs. **(A-F)** Cumulative distributions of log2 fold change (dominant negative / wildtype) for transcripts purified using rRNA depletion (ribo0) and poly(A) pulldown (pA) in zebrafish embryos depleted of individual mRNA decay factors: Dcp2 (A, D), Cnot7 (B, E), and both Dcp2 and Cnot7 (C, F) (black, transcripts with miR-430 target sites; red, transcripts with top 25% average codon pair half-life without miR-430 target sites; blue, transcripts with bottom 25% average codon pair half-life without miR-430 target sites; gray, remaining transcripts). Each p-value corresponds to the Kolmogorov-Smirnov test between the two distributions indicated by the colored dots. Note that Dcp2 shows differential decay on the ribo0, not the pA, suggesting Dcp2 can differentially degrade transcripts only after poly(A) tail removal. **(G-J)** Cumulative distributions of log2 fold change (upf1^-/-^/wildtype) for transcripts with top 25% long ORF (G) and bottom 25% short ORF (H), and maternal transcripts with top 25% long ORF (I) and bottom 25% short ORF (J) (black, transcripts with miR-430 target sites; red, transcripts with top 25% average codon pair half-life without miR-430 target sites; blue, transcripts with bottom 25% average codon pair half-life without miR-430 target sites; gray, remaining transcripts). Each p-value corresponds to the Kolmogorov-Smirnov test between the two distributions indicated by the colored dots. Note that the long ORF shows differential codon pair stability upon knockout of Upf1, whereas the short ORF shows a much weaker difference.

## References

1. Radhakrishnan, A., and Green, R. (2016). Connections Underlying Translation and mRNA Stability. J Mol Biol 428, 3558–3564. 10.1016/j.jmb.2016.05.025.

2. Guhaniyogi, J., and Brewer, G. (2001). Regulation of mRNA stability in mammalian cells. Gene 265, 11–23. 10.1016/s0378-1119(01)00350-x.

3. Kojima, M., Hoppe, C., and Giráldez, A. (2024). The maternal-to-zygotic transition: reprogramming of the cytoplasm and nucleus. Nature reviews. Genetics 26, 245–267. 10.1038/s41576-024-00792-0.

4. Chang, H., Yeo, J., Kim, J.-g., Kim, H., Lim, J., Lee, M., Kim, H.H., Ohk, J., Jeon, H.- Y., Lee, H., et al. (2018). Terminal Uridylyltransferases Execute Programmed Clearance of Maternal Transcriptome in Vertebrate Embryos. Molecular Cell 70, 72–82.e77. 10.1016/j.molcel.2018.03.004.

5. Giraldez, A.J., Mishima, Y., Rihel, J., Grocock, R.J., Van Dongen, S., Inoue, K., Enright, A.J., and Schier, A.F. (2006). Zebrafish MiR-430 promotes deadenylation and clearance of maternal mRNAs. Science 312, 75–79. 10.1126/science.1122689.

6. Rouget, C., Papin, C., Boureux, A., Meunier, A.C., Franco, B., Robine, N., Lai, E.C., Pelisson, A., and Simonelig, M. (2010). Maternal mRNA deadenylation and decay by the piRNA pathway in the early Drosophila embryo. Nature 467, 1128–1132. 10.1038/nature09465.

7. Sha, Q.Q., Zhang, J., and Fan, H.Y. (2019). A story of birth and death: mRNA translation and clearance at the onset of maternal-to-zygotic transition in mammalsdagger. Biol Reprod 101, 579–590. 10.1093/biolre/ioz012.

8. Vejnar, C.E., Abdel Messih, M., Takacs, C.M., Yartseva, V., Oikonomou, P., Christiano, R., Stoeckius, M., Lau, S., Lee, M.T., Beaudoin, J.D., et al. (2019). Genome wide analysis of 3’ UTR sequence elements and proteins regulating mRNA stability during maternal-to-zygotic transition in zebrafish. Genome Res 29, 1100–1114. 10.1101/gr.245159.118.

9. Bazzini, A.A., Del Viso, F., Moreno-Mateos, M.A., Johnstone, T.G., Vejnar, C.E., Qin, Y., Yao, J., Khokha, M.K., and Giraldez, A.J. (2016). Codon identity regulates mRNA stability and translation efficiency during the maternal-to-zygotic transition. EMBO J 35, 2087–2103. 10.15252/embj.201694699.

10. Mishima, Y., and Tomari, Y. (2016). Codon Usage and 3’ UTR Length Determine Maternal mRNA Stability in Zebrafish. Molecular cell 61 *6*, 874–885. 10.1016/j.molcel.2016.02.027.

11. Musaev, D., Abdelmessih, M., Vejnar, C.E., Yartseva, V., Weiss, L.A., Strayer, E.C., Takacs, C.M., and Giraldez, A.J. (2024). UPF1 regulates mRNA stability by sensing poorly translated coding sequences. Cell Rep 43, 114074. 10.1016/j.celrep.2024.114074.

12. Presnyak, V., Alhusaini, N., Chen, Y.H., Martin, S., Morris, N., Kline, N., Olson, S., Weinberg, D., Baker, K.E., Graveley, B.R., and Coller, J. (2015). Codon optimality is a major determinant of mRNA stability. Cell 160, 1111–1124. 10.1016/j.cell.2015.02.029.

13. Buschauer, R., Matsuo, Y., Sugiyama, T., Chen, Y.H., Alhusaini, N., Sweet, T., Ikeuchi, K., Cheng, J., Matsuki, Y., Nobuta, R., et al. (2020). The Ccr4-Not complex monitors the translating ribosome for codon optimality. Science 368. 10.1126/science.aay6912.

14. Radhakrishnan, A., Chen, Y.H., Martin, S., Alhusaini, N., Green, R., and Coller, J. (2016). The DEAD-Box Protein Dhh1p Couples mRNA Decay and Translation by Monitoring Codon Optimality. Cell 167, 122–132 e129. 10.1016/j.cell.2016.08.053.

15. Webster, M.W., Stowell, J.A., and Passmore, L.A. (2019). RNA-binding proteins distinguish between similar sequence motifs to promote targeted deadenylation by Ccr4-Not. Elife 8. 10.7554/eLife.40670.

16. Hia, F., Wu, Y., Yoshinaga, M., Goto-Ito, S., Iwasaki, W., Imami, K., Toh, H., Han, P., Cai, T., Ohira, T., et al. (2026). Human DHX29 detects nonoptimal codon usage to regulate mRNA stability. Science 392, eadw0288. 10.1126/science.adw0288.

17. Zhu, X., Cruz, V.E., Zhang, H., Erzberger, J.P., and Mendell, J.T. (2024). Specific tRNAs promote mRNA decay by recruiting the CCR4-NOT complex to translating ribosomes. Science 386, eadq8587. 10.1126/science.adq8587.

18. Wu, Q., and Bazzini, A.A. (2023). Translation and mRNA Stability Control. Annu Rev Biochem 92, 227–245. 10.1146/annurev-biochem-052621-091808.

19. Alexaki, A., Kames, J., Holcomb, D.D., Athey, J., Santana-Quintero, L.V., Lam, P.V.N., Hamasaki-Katagiri, N., Osipova, E., Simonyan, V., Bar, H., et al. (2019). Codon and Codon-Pair Usage Tables (CoCoPUTs): Facilitating Genetic Variation Analyses and Recombinant Gene Design. J Mol Biol 431, 2434–2441. 10.1016/j.jmb.2019.04.021.

20. Gutman, G.A., and Hatfield, G.W. (1989). Nonrandom utilization of codon pairs in Escherichia coli. Proc Natl Acad Sci U S A 86, 3699–3703. 10.1073/pnas.86.10.3699.

21. Tats, A., Tenson, T., and Remm, M. (2008). Preferred and avoided codon pairs in three domains of life. BMC Genomics 9, 463. 10.1186/1471-2164-9-463.

22. Gamble, C.E., Brule, C.E., Dean, K.M., Fields, S., and Grayhack, E.J. (2016). Adjacent Codons Act in Concert to Modulate Translation Efficiency in Yeast. Cell 166, 679–690. 10.1016/j.cell.2016.05.070.

23. Forrest, M.E., Pinkard, O., Martin, S., Sweet, T.J., Hanson, G., and Coller, J. (2020). Codon and amino acid content are associated with mRNA stability in mammalian cells. PLoS One 15, e0228730. 10.1371/journal.pone.0228730.

24. Burke, P.C., Park, H., and Subramaniam, A.R. (2022). A nascent peptide code for translational control of mRNA stability in human cells. Nat Commun 13, 6829. 10.1038/s41467-022-34664-0.

25. Chen, K.Y., Park, H., and Subramaniam, A.R. (2024). Massively parallel identification of sequence motifs triggering ribosome-associated mRNA quality control. Nucleic Acids Res 52, 7171–7187. 10.1093/nar/gkae285.

26. Nirenberg, M.W., and Matthaei, J.H. (1961). The dependence of cell-free protein synthesis in E. coli upon naturally occurring or synthetic polyribonucleotides. Proc Natl Acad Sci U S A 47, 1588–1602. 10.1073/pnas.47.10.1588.

27. Rappol, T., Waldl, M., Chugunova, A., Hofacker, I.L., Pauli, A., and Vilardo, E. (2024). tRNA expression and modification landscapes, and their dynamics during zebrafish embryo development. Nucleic Acids Res 52, 10575–10594. 10.1093/nar/gkae595.

28. Beaudoin, J.-D., Novoa, E.M., Vejnar, C.E., Yartseva, V., Takacs, C.M., Kellis, M., and Giraldez, A.J. (2018). Analyses of mRNA structure dynamics identify embryonic gene regulatory programs. Nature Structural & Molecular Biology 25, 677–686. 10.1038/s41594-018-0091-z.

29. Takyar, S., Hickerson, R.P., and Noller, H.F. (2005). mRNA Helicase Activity of the Ribosome. Cell 120, 49–58. 10.1016/j.cell.2004.11.042.

30. Lundberg, S.M., and Lee, S.I. (2017). A Unified Approach to Interpreting Model Predictions. Advances in Neural Information Processing Systems 30 (NIPS 2017).

31. Bhat, P., Cabrera-Quio, L.E., Herzog, V.A., Fasching, N., Pauli, A., and Ameres, S.L. (2023). SLAMseq resolves the kinetics of maternal and zygotic gene expression during early zebrafish embryogenesis. Cell Reports 42. 10.1016/j.celrep.2023.112070.

32. Mishima, Y., and Tomari, Y. (2017). Pervasive yet nonuniform contributions of Dcp2 and Cnot7 to maternal mRNA clearance in zebrafish. Genes Cells 22, 670–678. 10.1111/gtc.12504.

33. Lawir, D.F., Sikora, K., O’Meara, C.P., Schorpp, M., and Boehm, T. (2020). Pervasive changes of mRNA splicing in upf1-deficient zebrafish identify rpl10a as a regulator of T cell development. Proc Natl Acad Sci U S A 117, 15799–15808. 10.1073/pnas.1917812117.

34. Ishibashi, K., Shichino, Y., Han, P., Wakabayashi, K., Mito, M., Inada, T., Kimura, S., Iwasaki, S., and Mishima, Y. (2024). Translation of zinc finger domains induces ribosome collision and Znf598-dependent mRNA decay in zebrafish. PLoS Biol 22, e3002887. 10.1371/journal.pbio.3002887.

35. Cabrera-Quio, L.E., Schleiffer, A., Mechtler, K., and Pauli, A. (2021). Zebrafish Ski7 tunes RNA levels during the oocyte-to-embryo transition. PLoS Genet 17, e1009390. 10.1371/journal.pgen.1009390.

36. Pinkard, O., McFarland, S., Sweet, T., and Coller, J. (2020). Quantitative tRNA-sequencing uncovers metazoan tissue-specific tRNA regulation. Nat Commun 11, 4104. 10.1038/s41467-020-17879-x.

37. Reimao-Pinto, M.M., Behrens, A., Forcelloni, S., Frohlich, K., Kaya, S., and Nedialkova, D.D. (2024). The dynamics and functional impact of tRNA repertoires during early embryogenesis in zebrafish. EMBO J 43, 5747–5779. 10.1038/s44318-024-00265-4.

38. Strayer, E.C., Krishna, S., Lee, H., Vejnar, C., Neuenkirchen, N., Gupta, A., Beaudoin, J.D., and Giraldez, A.J. (2024). NaP-TRAP reveals the regulatory grammar in 5’UTR-mediated translation regulation during zebrafish development. Nat Commun 15, 10898. 10.1038/s41467-024-55274-y.

39. Absmeier, E., Chandrasekaran, V., O’Reilly, F.J., Stowell, J.A.W., Rappsilber, J., and Passmore, L.A. (2023). Specific recognition and ubiquitination of translating ribosomes by mammalian CCR4-NOT. Nat Struct Mol Biol 30, 1314–1322. 10.1038/s41594-023-01075-8.

40. Bae, H., and Coller, J. (2022). Codon optimality-mediated mRNA degradation: Linking translational elongation to mRNA stability. Mol Cell 82, 1467–1476. 10.1016/j.molcel.2022.03.032.

41. Zhang, H., Zhang, L., Lin, A., Xu, C., Li, Z., Liu, K., Liu, B., Ma, X., Zhao, F., Jiang, H., et al. (2023). Algorithm for optimized mRNA design improves stability and immunogenicity. Nature 621, 396–403. 10.1038/s41586-023-06127-z.

42. Jia, L., Mao, Y., Ji, Q., Dersh, D., Yewdell, J.W., and Qian, S.B. (2020). Decoding mRNA translatability and stability from the 5’ UTR. Nat Struct Mol Biol 27, 814–821. 10.1038/s41594-020-0465-x.

43. Rahaman, S., Faravelli, S., Voegeli, S., and Becskei, A. (2023). Polysome propensity and tunable thresholds in coding sequence length enable differential mRNA stability. Sci Adv 9, eadh9545. 10.1126/sciadv.adh9545.

44. Garzia, A., Jafarnejad, S.M., Meyer, C., Chapat, C., Gogakos, T., Morozov, P., Amiri, M., Shapiro, M., Molina, H., Tuschl, T., and Sonenberg, N. (2017). The E3 ubiquitin ligase and RNA-binding protein ZNF598 orchestrates ribosome quality control of premature polyadenylated mRNAs. Nat Commun 8, 16056. 10.1038/ncomms16056.

45. Burow, D.A., Martin, S., Quail, J.F., Alhusaini, N., Coller, J., and Cleary, M.D. (2018). Attenuated Codon Optimality Contributes to Neural-Specific mRNA Decay in Drosophila. Cell Reports 24, 1704–1712. 10.1016/j.celrep.2018.07.039.

46. Vejnar, C.E., and Giraldez, A.J. (2020). LabxDB: versatile databases for genomic sequencing and lab management. Bioinformatics 36, 4530–4531. 10.1093/bioinformatics/btaa557.

47. Langmead, B., and Salzberg, S.L. (2012). Fast gapped-read alignment with Bowtie 2. Nat Methods 9, 357–359. 10.1038/nmeth.1923.

48. Virtanen, P., Gommers, R., Oliphant, T.E., Haberland, M., Reddy, T., Cournapeau, D., Burovski, E., Peterson, P., Weckesser, W., Bright, J., et al. (2020). SciPy 1.0: fundamental algorithms for scientific computing in Python. Nature Methods 17, 261–272. 10.1038/s41592-019-0686-2.

49. Harris, C.R., Millman, K.J., van der Walt, S.J., Gommers, R., Virtanen, P., Cournapeau, D., Wieser, E., Taylor, J., Berg, S., Smith, N.J., et al. (2020). Array programming with NumPy. Nature 585, 357–362. 10.1038/s41586-020-2649-2.

50. McKinney, W. (2010). Data Structures for Statistical Computing in Python. Proceedings of the 9th Python in Science Conference.

51. Hunter, J.D. (2007). Matplotlib: A 2D graphics environment. Computing in Science & Engineering 9, 90–95. 10.1109/MCSE.2007.55.

52. Pedregosa, F., Varoquaux, G., Gramfort, A., Michel, V., Thirion, B., Grisel, O., Blondel, M., Prettenhofer, P., Weiss, R., Dubourg, V., et al. (2011). Scikit-learn: Machine Learning in Python. Journal of Machine Learning Research 12, 2825–2830.

53. Chen, T., and Guestrin, C. (2016). XGBoost: A Scalable Tree Boosting System. Proceedings of the 22nd ACM SIGKDD International Conference on Knowledge Discovery and Data Mining.

54. Lorenz, R., Bernhart, S.H., Höner zu Siederdissen, C., Tafer, H., Flamm, C., Stadler, P.F., and Hofacker, I.L. (2011). ViennaRNA Package 2.0. Algorithms for Molecular Biology 6. 10.1186/1748-7188-6-26.

55. Leppek, K., Schott, J., Reitter, S., Poetz, F., Hammond, M.C., and Stoecklin, G. (2013). Roquin promotes constitutive mRNA decay via a conserved class of stem-loop recognition motifs. Cell 153, 869–881. 10.1016/j.cell.2013.04.016.

56. Kyte, J., and Doolittle, R.F. (1982). A simple method for displaying the hydropathic character of a protein. J Mol Biol 157, 105–132. 10.1016/0022-2836(82)90515-0.

57. Grimsley, G.R., Scholtz, J.M., and Pace, C.N. (2009). A summary of the measured pK values of the ionizable groups in folded proteins. Protein Sci 18, 247–251. 10.1002/pro.19.

58. Prohaska, T., Irrgeher, J., Benefield, J., Böhlke, J.K., Chesson, L.A., Coplen, T.B., Ding, T., Dunn, P.J.H., Gröning, M., Holden, N.E., et al. (2022). Standard atomic weights of the elements 2021 (IUPAC Technical Report). Pure and Applied Chemistry 94, 573–600. 10.1515/pac-2019-0603.

59. Huang, Y., Lin, T., Lu, L., Cai, F., Lin, J., Jiang, Y.E., and Lin, Y. (2021). Codon pair optimization (CPO): a software tool for synthetic gene design based on codon pair bias to improve the expression of recombinant proteins in Pichia pastoris. Microb Cell Fact 20, 209. 10.1186/s12934-021-01696-y.

